# The atypical RHO-GTPase RND3/RHOE interacts with FLRT3 to regulate cortical migration and folding

**DOI:** 10.64898/2026.07.10.737444

**Authors:** B Zammou, P Marfull-Oromí, S Aghrabatt, N Nicholson-Sabaté, M Pàmpols, Vilà Ll, M Berbeira-Santana, V Fernández, D Nagerl, D Chauhan, X Dolcet, M Encinas, R Klein, V Borrell, E Seiradake, D del Toro, J Egea

## Abstract

Cortical gyrification, characterized by the formation of sulcal fissures and gyral peaks, is a distinctive feature of large mammalian brains and is associated with the emergence of higher cognitive functions. This contrasts with small mammals, like mice, which generally display smooth (lissencephalic) cortices. Gyrification results from the integration of several complex processes such as neuron progenitor amplification and migration. However, the underlying molecular mechanisms remain poorly understood. Here, we show that genetic ablation of *Rnd3*, encoding the atypical RHO-GTPase RND3/RHOE, induces spontaneous cortical sulci in a subset of mouse embryos without increasing neural progenitor amplification. Instead, RND3 regulates cortical neuron migration, as RND3 overexpression delays neuronal migration, whereas its loss accelerates this process. Similar phenotypes reported in the double *Flrt1*/*3* mutants, suggested a functional interaction between the transmembrane FLRTs and RND3 during cortex development. We demonstrate that *Rnd3* and *Flrt3* are co-expressed in migrating cortical neurons and interact through a β-strand-mediated binding mechanism, involving a conserved motif in RND3. Disruption of this motif abolishes FLRT3 binding, impairs RND3-mediated regulation of neuronal migration and promotes sulcus formation, indicating that FLRT3-RND3 suppresses cortical folding by regulating neuronal migration in mice. Consistently, *Rnd3* expression is reduced in the outer subventricular zone of the prospective sulcal regions in the gyrencephalic ferret brain, supporting the notion that RND3 downregulation contributes to sulci development. Together, these findings identify FLRT3-RND3 signalling as a conserved negative regulator of cortical neuron migration and cortical folding.

## Introduction

The neocortex is the youngest part of the cerebral cortex and the brain region undergoing the largest expansion during evolution. Remarkably, in gyrencephalic species like humans, the neocortex is highly folded forming peaks (gyri) and fissures (sulci). This trait was an important evolutionary feature in the acquisition of complex cognitive functions in large mammals, allowing the increase of cortical surface area in a reduced cranial volume and facilitating the interconnectivity of distal cortical regions^1,2,3^. In contrast, mice and other smaller mammals display a smooth cortex with no folds, and are therefore classified as lissencephalic. Cortical folding occurs mainly during embryonic development and early postnatal stages, and errors of this process underly severe human pathologies such as polymicrogyria, cortical dysplasia, pachygyria or lissencephaly^4,5,6,7^. Functional studies in mice and gyrencephalic animal models such as the ferret have revealed some of the genetic and cellular mechanisms for cortex expansion and folding^8,9^. Among these mechanisms, the emergence of a specific subdivision of the subventricular zone (SVZ) in gyrencephalic species, the outer SVZ (OSVZ), played a key evolutionary role by expanding the pool of basal progenitors (BPs), which thereby supported the increased neuronal production necessary for cortical tangential expansion^10,11,12,13,14,15,16,17^. Importantly, different regional cortical growth rate, specially within the OSVZ, has been shown to correlate with the prospective formation of gyri (higher proliferation) and sulci (lower proliferation)^14^. In addition, neuron migration also plays a crucial role during the development of cortical folds. In gyrencephalic species, cortical neurons follow divergent and more sinuous pathways, instead of the parallel trajectories found in lissencephalic brains, allowing tangential expansion which generates mechanical tension and cortical folding^3,14,15,18,19^. Finally, cortical folding has been shown to be regulated by cell adhesion, involving components of the extracellular matrix as well as cell-surface adhesion receptors^20^.

FLRTs (FLRT1-3, <u>F</u>ibronectin and <u>L</u>eucine <u>R</u>ich <u>T</u>ransmembrane proteins) are single-pass transmembrane proteins including a short (∼100 amino acids), conserved C-terminal intracellular domain (ICD)^21^. They display an extraordinary functional versatility during cortex development with diverse mechanisms of action, both cell autonomous and non-cell autonomous, as chemorepulsive signals, co-receptors or cell adhesion molecules^22^. FLRTs act as chemorepellent ligands for the netrin receptors Unc5 modulating radial migration of BPs and the distribution of cortical interneurons^23,24^. In addition, FLRT3 is required for the fine tuning of the topographic arrangement of thalamocortical axons acting as a co-receptor of the Slit1 receptor Robo1^25^. Remarkably, double genetic ablation of *Flrt1* and *Flrt3* in mice, causes the appearance of spontaneous cortical sulci during development due to altered neuron migration^26^. Interestingly, combining this model with genetic lines that induce progenitor expansion greatly enhances cortical folding, indicating that migration and proliferation act synergistically on cortical folding^27^. Notably, in the gyrencephalic ferret brain *Flrt1* and *Flrt3* expression are reduced in the OSVZ of areas of prospective sulcal development, strongly supporting the importance of FLRTs for cortical folding in gyrencephalic species^26^. Mechanistically, deletion of *Flrt1/3* causes a reduction of intercellular adhesion that leads to faster neuronal migration and to neuronal clustering in the cortical plate (CP), due to the action of FLRTs as homotypic or heterotypic adhesion molecules to Latrophilins^28,29,30^. The intracellular signalling mechanisms triggered by these adhesive properties of FLRTs during cortex development leading cortical folding have not been investigated.

RND3 (RHOE/RHO8) belongs to a small family of atypical RHO GTPases including RND1 (RHO6) and RND2 (RHON/RHO7)^31^. Like other RHO-family members, RND proteins control actin cytoskeleton dynamics^32^. In contrast, RNDs are always in a GTP-bound state due to the lack of GTPase activity and are consequently considered constitutive active proteins^31,32,33^. The regulation of RND activity depends on mechanisms different from the GDP/GTP exchange such as gene transcription, protein stability, and post-translational modifications including site-specific phosphorylations or farnesylation, which tether these proteins to the plasma membrane^34^. RND3 control of cytoskeleton dynamics is important for cell morphology and migration in fibroblasts, cancer cells and neurons, by antagonizing the RHOA signaling pathway^31,35,36,37,38,39,40^. *Rnd3* KO mice exhibit early aqueductal stenosis and subsequent hydrocephaly and the complete lack of the perineal nerve^41,42^. In neurons, RND3 is necessary for locomotion phase in the cortical plate (CP) during radial neuron migration in the cortex, interkinetic nuclear migration of radial glial stem cells, proliferation of BPs, neuronal polarization, oligodendrocyte differentiation, SVZ and rostral migratory stream formation, and for the maintenance of the radial fiber scaffold in the subpallium^43,44,45,46,47,48,49,50^. In addition, RND3 is required for the correct development of thalamocortical and striatal projections and of subpallial structures derived from the medial ganglionic eminence, such as the globus pallidus^51^. Interestingly, the role of RND3 in cortical neuron migration is finely regulated through its interaction with transmembrane receptors such as PlexinB2, enabling cell-extrinsic signals mediated by these receptors to modulate RND3 activity.^50,51^. On the other hand, in the *Xenopus* embryo, RND1 interaction with FLRT3 acts downstream of TGFβ signaling to control cell adhesion during gastrulation^54,55^. However, the role of this FLRT-RND interaction for the formation of the nervous system is currently unknown.

Here we show that genetic ablation of *Rnd3* induces spontaneous cortical sulci in a fraction of KO embryos displaying features that resemble genuine sulci of gyrencephalic species. In these mutant embryos, neuron proliferation remains largely unaltered but cortical radial migration is affected. Thus, in utero electroporation (IUE) experiments show that RND3 overexpression delays neuronal migration, whereas RND3 ablation accelerates it; an effect being regulated by RND3 phosphorylation and its localization at the cell membrane. Notably, these phenotypes are reminiscent of those of *Flrt1/3* double KOs (DKOs)^26^, suggesting a functional link between RND3 and FLRTs during cortical neuron migration. Indeed, we show that *Rnd3* and *Flrt3* are co-expressed in migrating cortical neurons during early mouse development, we then demonstrate that RND3 and FLRT3 interact and identify the FLRT3-binding motif (F3BM) in RND3 using AlphaFold 2.0. We found that mutation of this motif (RND3^F3BM^) abolishes the migration effects of wild-type RND3, and induces cortical sulci. These findings indicate that FLRT3-RND3 signalling regulates cortical radial migration and suppresses cortical folding in mice. In full agreement with this notion, *Rnd3* expression in the OSVZ of normally developing ferrets was significantly reduced in areas of prospective sulci development. Together, we propose the interaction FLRT3-RND3 as a key regulatory mechanism of cortical development and folding.

## Methods

### Mouse lines and ferret

Mouse care was performed in accordance with the Guidelines of University of Lleida for Animal Experimentation in accordance wit h Catalan, Spanish, and European Union regulations (Decret 214/1997, Real Decreto 53/2013 and Directive 63/2010). Animals were housed in the animal house of the University of Lleida with 12:12h light/dark cycle and food/water available ad libitum. The knock-out alleles *Rnd3^gt^* (<u>g</u>ene trap allele with a lacZ cassette), the *Rnd3^tm1d^*, the conditional allele *Rnd3^tm1c^* (or *Rnd3^lx^*) and the Nestin-Cre mouse line were previously described^51^. The *Rnd3^tm1d^* allele was obtained by crossing *Rnd3^tm1c/tm1c^*(*Rnd3^lx/lx^*) females with males carrying the deleter CAG-CRE line (B6.Cg-Tg(CAG-Cre)CZ-M02Osb, RBRC01828, RIKEN BRC). Littermates with genotypes *Rnd3^+/+^*, *Rnd3^gt/+^*, *Rnd3^tm1d/+^* were used as controls. Transgenic mice were backcrossed with C57BL/6. The day of positive vaginal plug was considered embryonic (E) day 0.5 and pregnant females were sacrificed by quick cervical dislocation at different embryonic days, as indicated. Embryos were sacrificed by decapitation.

Sable pigmented ferrets (*Mustela putorius furo*) were obtained from Euroferret and kept on a 16:8h light:dark cycle at the Animal Facilities of the Universidad Miguel Hernández. Ferrets were analysed independently of their gender. All animals were treated according to Spanish and EU regulations, and experimental protocols were approved by the Universidad Miguel Hernández Institutional Animal Care and Use Committee (IACUC).

### βGAL staining

Embryonic brains were fixed for 4 hr in 4% paraformaldehyde (PFA)/PBS, cryoprotected in 30% sucrose in PBS, embedded in cryoprotective Tissue-Freezing Medium (Electron Microscopy Sciences, 72592), and stored at −80°C. Serial 20 μm coronal sections were made in cryostat (Leica CM3000 or CM1950) and collected in Superfrost Plus™ slides (Thermo Fisher). Cryosections of Rnd3gt/+ mice were washed and stained with XGal staining solution at 22°C some hours until blue precipitate was observed. Finally, slices were washed and mounted with Fluoromount-G (SouthernBiotech, Cat. No.0100–01). If co-immunostaining was desired, slides were washed with PBS and continued with normal immunofluorescence protocol. Images were obtained with a transmitted light microscope (Olympus BX51).

### Tissue immunofluorescence, microscopy and image analysis

Embryonic brains and slices were obtained as indicated before. Sections were washed with PBS, permeabilized, and blocked with 5% donkey serum in 0.1% Triton X-100 in PBS for 1 hr at 22°C. Then, slides were incubated with the following primary antibodies diluted in blocking buffer for 12–24 hr at 4°C: anti-CTIP2 (Abcam, ab-18465), anti-TBR1 (Abcam, ab31940), anti-SATB2 (Abcam, ab51502), anti-laminin (Abcam, ab11575), anti-PH3 (Abcam, ab5176), anti-ISLR2 (R&D, AF4650), anti-GFP (Rockland, 600-101-215) and anti-βGal (Cappel, 559761). After washing, samples we incubated with fluorescent secondary antibodies (1:300, Jackson ImmunoResearch) diluted in blocking buffer containing DAPI (1:1,000). Samples were mounted with antifading mounting medium Fluoromount-G (Southern Biotech) and visualized with an upright fluorescence microscope (Olympus BX51) or with a confocal laser scanning biological microscope FV1000, FLUOVIEW (Olympus). Brightness and contrast were linearly adjusted with Image J. For quantification, images were classified into rostral (before the emergence of the thalamus), intermediate (until thalamocortical axons cross the diencephalic-telencephalic boundary) and caudal (until the end of the cortex). Staining were counted manually or with the help of CellProfiler (Broad Institute^56^) and the CellCounter plugin in Image J. Curvature analysis of the radial glial processes was calculated using the Kappa plugin for ImageJ^57^.

### Expression vectors

Mouse *Rnd3* cDNA containing a N-terminus FLAG tag was kindly provided by Dr. Ignacio Pérez-Roger (Universidad CEU Cardenal Herrera, Valencia, Spain). Mouse full-length FLRT3 and the mutant where the entire intracellular domain is substituted by EGFP (FLRT3^ΔCEGFP^) were previously described^25^. Constructs were subcloned into the mammalian expression vectors pcDNA3.1 (ThermoFisher), for HEK293T cell transfection), or pCAGIG (Addgene, 11159), for IUE, which contains an internal ribosome entry site (IRES) and the EGFP as a reporter gene for tracking the electroporated cells.

### Generation of RND3 mutant constructs

To generate the different RND3 mutants, we subcloned the mouse *Rnd3* cDNA (N-terminus FLAG tagged) into pBluescript. Site-directed mutagenesis to substitute S210, S218 and S240 by alanines and generate RND3^P1P2P3^ mutant (sequentially mutating P1→P2→P3) or to substitute the FENYTASF sequence by PGTPTPP and generate the RND3^F3BM^ mutant were performed by PCR (AccuPrime™ Pfx SuperMix, ThermoFisher) the whole plasmid with primers containing the desired mutations (indicated in bold and underlined): P1Fw, CGTTAAGCGGAACAAA**<u>G</u>**CGCAGAGGGCCACAAAGCG and P1Rv, CGCTTTGTGGCCCTCTGCG**<u>C</u>**TTTGTTCCGCTTAACG; P2Fw, GCCACAAAGCGGATT**<u>G</u>**CGCACATGCCTAGCAGACC and P2Rv, GGTCTGCTAGGCATGTGCG**<u>C</u>**AATCCGCTTTGTGGC; P3Fw, CGAAAGGACAAAGCGAAG**<u>GC</u>**CTGTACTGTGATGTGAAAGCTATCGC and P3Rv: GCGATAGCTTTCACATCACAGTACAG**GC**CTTCGCTTTGTCCTTTCG; F3BMFw, CCCAGAAAATTACGTCCCTACGGTG**<u>CC</u>**TG**<u>G</u>**GA**<u>C</u>**T**<u>CC</u>**CACT**<u>C</u>**CC**<u>CC</u>**T**<u>CC</u>**TGAAATC GACACACAAAGAATAGAGTTGAGCC and F3BMRv, GGCTCAACTCTATTCTTTGTGTGTCGATTTCA**<u>GG</u>**A**<u>GG</u>**GG**<u>G</u>**AGTG**<u>GG</u>**A**<u>G</u>**TC**<u>C</u>**CA**<u>GG</u>** CACCGTAGGGACGTAATTTTCTGGG. Generation of the RND3^ΔFS^ was achieved by PCR-amplifying (AccuPrime™ Pfx SuperMix, ThermoFisher) the entire *Rnd3* cDNA including a reverse primer containing a premature STOP codon just before the CAAX motif at the C-terminus end: FSFw, CCC*<u>CTCGAG</u>***ATG<u>GACTACAAGGACGACGATGACAAG</u>**AATTCG (*<u>XhoI</u>*, **Start Codon**, **<u>FLAG</u>**) and FSRv, CCC*<u>GCGGCCGC</u>*CTT**<u>TCA</u>**GCTCTTCGCTTTGTCCTTTCGTAAGTCCG (*NotI* and new **<u>Stop Codon</u>**). All the mutations were verified by restriction analysis and sequencing. The mutated *Rnd3* cDNAs were subcloned as above into the mammalian expression vector pcDNA3.1 or pCAGIG.

### Co-immunoprecipitation, Western-blot and immunofluorescence in HEK293T cells

For co-immunoprecipitation and Western-blot assays, HEK293T cells were transfected by a home-made calcium phosphate method with the following constructs: full-length FLRT3, FLRT3^ΔCEGFP^, full-length RND3, RND3^P1P2P3^, RND3^F3BM^ or FLRT3^ΔFS^ (all RND3 constructs are N-terminus FLAG tagged). After 36 hours the cells were lysed in lysis buffer (150mM NaCl, 20mM Tris [pH7.4], 10% glycerol, 1% Triton X-100, 1mM Na3VO4, 1mM EDTA, 5mM NaPP, 5mM NaF and a protease inhibitor cocktail [Complete EDTA-free from Roche]) and cleared by centrifugation. Lysates were directly analysed by Western-blot (total cell lysates, TCL) or incubated for 4 hours at 4°C on a rotating wheel with the ANTI-FLAG® M2 Affinity Gel resin (Sigma). TCL and immunoprecipitates were resolved by SDS-PAGE, transferred to PVDF membranes (Millipore), and analysed by Western-blot with primary antibodies against FLAG (anti-FLAG antibody, Sigma-Aldrich, F7425) or FLRT3 (extracellular domain epitope, R&D AF2795) and the appropriate HRP secondary antibodies (Jackson Immunoresearch). Bands were revealed using the chemiluminescent HRP substrate Immobilon Western (Millipore). Images of the blots and quantification of band intensity were obtained using the ImageLab software (BioRad) subtracting the background signal in each line. Quantifications were expressed as fold signal over the non-transfected cells. For immunofluorescence, HEK293T cells were seeded in poly-D-lysine (Sigma-Andrich, P7886)-coated glass coverslips, transfected as above and, after 24h, fixed with 4% PFA+4% sucrose in PBS, permeabilized with 0.1% Triton X-100 in PBS and blocked with 4% donkey serum in PBS. Primary antibody against FLAG (anti-FLAG antibody, Sigma-Aldrich, F7425) and the appropriate secondary antibody (Jackson Immunoresearch), were used. Images were taken with a confocal laser scanning biological microscope FV1000, FLUOVIEW (Olympus).

### Intraventricular injection and IUE

E13.5-E15.5 pregnant C57BL/6 wild-type mice were deeply anesthetized with isoflurane (IsoFlo, Zoetis) during the entire operation procedure. To relax uterus muscles, β2 agonist Ritodrine (Sigma R0758) was administered intraperitoneally, and buprenorphine (Buprex, 100 mg/ml) subcutaneously as an analgesic. A 2 cm laparotomy section was made in the abdomen, and the uterine horns were carefully exposed and lubricated with NaCl 0.9% at 37° C. Two to 4 microliters of purified plasmid DNA dissolved in PBS (1 μg/μl) containing 0.025% of Fast Green (Sigma-Aldrich) was injected in the lateral ventricles of each embryo using a glass capillary (World Precision Instruments) sharpened previously by Puller P-97 (Sutter Instrument). Platinum electrodes (CUY701P20L, Nepagene) were placed across the head with the positive pole adjacent to the neocortex, to enhance the permeability of the cell membrane and allow the entrance of DNA. Five 30 mV electric pulses of 50 ms with intervals of 950 ms were charged by an electroporator (ECM830, BTX). Uterine horns were placed back into the abdominal cavity and abdomen wall and skin were surgically sutured. During the whole operation embryos were manipulated with ring forceps (Fine Science Tools). Embryonic brains were dissected, as indicated, at different time-points after electroporation and fixed directly in ice cold 4% paraformaldehyde (PFA) solution in PBS overnight.

### AlphaFold analysis and protein sequence anaylisis

We used AlphaFold3^58^ for the structural prediction of FLRT1-3/RND1-3 complexes, using FLRT intracellular domain (ICD) sequences. TMHMM2.0 (https://services.healthtech.dtu.dk/services/TMHMM-2.0/) was used to predict the N-terminal ICD boundary. Briefly, the sequences used are: mouse FLRT1 (NP958813.1) residues 579-674 (RAGELLTRERVYNRGSRRKDDYMESGTKKDNSILEIRGPGLQMLPINPYRSKEEYVVHTIFPSNGSSLCKGAHTIGYGTTRGYREAGIPDVDYSYT), mouse FLRT2 (NP958926.1) residues 564-660 (HMHKKGRYTSQKWKYNRGRRKDDYCEAGTKKDNSILEMTETSFQIVSLNNDQLLKGDFRLQPIYTPNGGINYTDCHIPNNMRYCNSSVPDLEHCHT), mouse FLRT3 (NP001165631.1) residues 552-649 (HRNGSLFSRNCAYSKGRRRKDDYAEAGTKKDNSILEIRETSFQMLPISNEPISKEEFVIHTIFPPNGMNLYKNNLSESSSNRSYRDSGIPDSDHSHS), mouse RND1 (NP766200.1) residues 1-232 (MKERRAPQPVVVRCKLVLVGDVQCGKTAMLQVLAKDCYPETYVPTVFENYTACLETEEQRVELSLWDTSGSPYYDNVRPLCYSDSDAVLLCFDISRPETMDSALKKWRTEILDYCPSTRVLLIGCKTDLRTDLSTLMELSHQKQAPISYEQGCAIAKQLGAEIYLEGSAFTSETSIHSIFRTASMVCLNKSSPVPPKSPVRSLSKRLLHLPSRSELISTTFKKEKAKSCSIM), mouse RND2 (NP033838.1) residues 1-227 (MEGQSGRCKIVVVGDAECGKTALLQVFAKDAYPGSYVPTVFENYTASFEIDKRRIELNMWDTSGSSYYDNVRPLAYPDSDAVLICFDISRPETLDSVLKKWQGETQEFCPNAKVV LVGCKLDMRTDLATLRELSKQRLIPVTHEQGTVLAKQVGAVSYVECSSRSSERSVRDVFHVATVASLGRGHRQLRRTDSRRGLQRSTQLSGRPDRGNEGEMHKDRAKSCNLM), mouse RND3 (NP083086.1) residues 1-244 (MKERRASQKLSSKSIMDPNQNVKCKIVVVGDSQCGKTALLHVFAKDCFPENYVPTVFENYTASFEIDTQRIELSLWDTSGSPYYDNVRPLSYPDSDAVLICFDISRPETLDSVLKKWKGEIQEFCPNTKMLLVGCKSDLRTDVSTLVELSNHRQTPVSYDQGANMAKQIGAATYIECSALQSENSVRDIFHVATLACVNKTNKNVKRNKSQRATKRISHMPSRPELSAVATDLRKDKAKSCTVM)

We then identified the predicted FLRT1-3/RND1-3 interfaces using PDBePISA^59^ and calculated the times each interacting amino acid is present for each RND isoform. We utilized the COUNTIFS formula in Excel to give a score from 1-3 for each amino acid (1 meaning it appears in one RND/FLRT1-3 complex and 3 meaning it appears for all three RND-FLRT1-3 complexes) and created a heatmap onto the Rnd1-3 wild type predicted structures in pymol.

For the multiple sequence alignments of FLRT1-3 and RND1-3 sequences we utilized Clustal Omega, and the following Uniprot sequences: mouse FLRT1 (Q6RKD8), human FLRT1 (Q9NZU1), rat FLRT1 (D4AC39), chicken FLRT1 (A0A8V0X339), bovine FLRT1 (G3MY74), xenopus FLRT1 (Q6DJD2), zebrafish FLRT1 (A0A8M9PLL4/A0AC58H8M3), mouse FLRT2 (Q8BLU0), human FLRT2 (O43155), rat FLRT2 (D3ZTV3), chicken FLRT2 (E1C6W5), bovine FLRT2 (F1MVT1), xenopus FLRT2 (A0A1L8F9F1), zebrafish FLRT2 (A0A8M1NFA3), mouse FLRT3 (Q8BGT1), human FLRT3 (Q9NZU0), rat FLRT3 (B1H234), chicken FLRT3 (F1NUK7), bovine FLRT3 (F1N0R7), xenopus FLRT3 (Q70AK3), zebrafish FLRT3 (B8A507), mouse RND1 (Q8BLR7), human RND1 (Q92730), rat RND1 (Q5FVG9), bovine RND1 (Q2HJ68), xenopus RND1 (Q9W761), zebrafish RND1 (Q6TEM4), mouse RND2 (Q9QYM5), human RND2 (P52198), rat RND2 (Q5HZE6), chicken RND2 (F1N9W4), bovine RND2 (A0AAA9TJS7), zebrafish RND2 (Q0P4B0), mouse RND3 (P61588), human RND3 (P61587), rat RND3 (Q6SA80), chicken RND3 (A0A8V0XYC4), bovine RND3 (A5PKJ7), xenopus RND3 (Q6DJ10), and zebrafish RND3 (Q6PC71).

We used AlphaFold2^60^ for the structural prediction of RND1-3 wild type. We used the following sequences: mouse RND1 (NP766200.1) residues 1-232 (MKERRAPQPVVVRCKLVLVGDVQCGKTAMLQVLAKDCYPETYVPTVFENYTACLETEEQRVELSLWDTSGSPYYDNVRPLCYSDSDAVLLCFDISRPETMDSALKKWRTEILDYCPSTRVLLIGCKTDLRTDLSTLMELSHQKQAPISYEQGCAIAKQLGAEIYLEGSAFTSETSIHSIFRTASMVCLNKSSPVPPKSPVRSLSKRLLHLPSRSELISTTFKKEKAKSCSIM),mouse RND2 (NP033838.1) residues 1-227 (MEGQSGRCKIVVVGDAECGKTALLQVFAKDAYPGSYVPTVFENYTASFEIDKRRIELNMWDTSGSSYYDNVRPLAYPDSDAVLICFDISRPETLDSVLKKWQGETQEFCPNAKVVLVGCKLDMRTDLATLRELSKQRLIPVTHEQGTVLAKQVGAVSYVECSSRSSERSVRDVFHVATVASLGRGHRQLRRTDSRRGLQRSTQLSGRPDRGNEGEMHKDRAKSCNLM), mouse RND3 (NP083086.1) residues 1-244 (MKERRASQKLSSKSIMDPNQNVKCKIVVVGDSQCGKTALLHVFAKDCFPENYVPTVFENYTASFEIDTQRIELSLWDTSGSPYYDNVRPLSYPDSDAVLICFDISRPETLDSVLKKWKGEIQEFCPNTKMLLVGCKSDLRTDVSTLVELSNHRQTPVSYDQGANMAKQIGAATYIECSALQSENSVRDIFHVATLACVNKTNKNVKRNKSQRATKRISHMPSRPELSAVATDLRKDKAKSCTVM). For the Rnd3 F3BM mutant we substituted the FENYTASF sequence by PGTPTPP.

Structural models were visualized in ChimeraX^61^ and Pymol (The PyMOL Molecular Graphics System, Version 3.0 Schrödinger, LLC). The top ranked solution (highest confidence score) is shown.

### ConSurf and conservation analysis

To produce the conservation scores for the structures we used ConSurf Colab (https://colab.research.google.com/drive/1PhDXX7k12oUsV6T_xkXC3Rm9R99e7tHz# scrollTo=zBTwgYcIK6Ul)^62^. For the ConSurf run we used the standard settings, but we inputted a custom MSA using Clustal Omega and the RND3 wild type structural prediction. To generate the custom MSA, we used the following sequences for RND1 from Uniprot: zebrafish (Q6TEM4), pheasant (A0A669QEH4), mouse (Q8BLR7), snake (AOA670YM58), caecilian (A0A6P8R289), the following for RND2: zebrafish (Q0P4B0), pheasant (AOA669PRZ8), mouse (Q9QYM5), snake (A0A670ZN47), caecilian (A0A6P8PFX6); and the following for RND3: zebrafish (Q6PC71), pheasant (A0A669PG61), mouse (P61588), snake (A0A670ZCN7), caecilian (A0A6P8QXI2). The conservation scores were mapped onto the structures using Pymol and the ConSurf provided script.

### In situ hybridization with ferret *Rnd3* probe and quantification of the signal

For in situ hybridization of *Rnd3* in ferret brains, a new ferret probe was designed based on *Rnd3* cDNA sequence retrieved from Ensembl.org database (transcript ENSMPUT00000000333.1). The primers for cloning the probe by PCR were: *Rnd3^MP^*-forward, CCC<u>CTCGAG</u>CCCCGTCTGCGCGGCGCTTCCAGAGTTCCC (containing a 5’ end <u>XhoI</u> restriction site) and *Rnd3^MP^*-reverse, CCC<u>GAATTC</u>AATGTGACTCTGGCTGTGCACGTCACTCGG (containing a 5’ end <u>EcoRI</u> restriction site). These primers amplify a 981 bp cDNA fragment containing the entire 735 bp of the ferret *Rnd3* coding sequence, 138 bp of the 5’ UTR and 108 bp of the 3’ UTR. Early ferret postnatal brains (P4-6) were homogenized with TRIzol (ThermoFisher) for total RNA extraction and cDNA synthesis was performed with random primers (Protoscript II, New England Biolabs). PCR with *Rnd3^MP^*-forward and *Rnd3^MP^*-reverse primers was carried out with AccuPrime™ Pfx SuperMix (ThermoFisher) and the PCR product was subcloned into TOPO2.1 vector (ThermoFisher), according to manufacturer’s instructions. Correct cloning of the ferret *Rnd3* cDNA was verified by restriction analysis and sequencing. Sense and anti-sense *RNA* probes were synthesized and labelled with digoxigenin (DIG; Roche Diagnostics) according to the manufacturer’s instructions. In situ hybridization in ferret sections was performed as described previously^26^. Quantification of the signal intensity of *Rnd3* in situ hybridization on ferret brain sections was performed by using ImageJ software. Images were first converted to 32-bit and inverted before further processing. For the generation of *Rnd3* stain intensity plots along the rostro-caudal axis of the ferret brain the entire thickness of the OSVZ was tracked with a segmented line which was straightened using the straighten plugin in ImageJ (http://rsbweb.nih.gov/ij/plugins/straighten.html). Then, pixel intensity along the length of the OSVZ was analyzed using Plot Profile and raw data was smoothened to better visualized the intensity trend. To compare *Rnd3* stain intensity between sulcus and the adjacent gyrus in different cortical layers (CP, OSVZ or ISVZ), rectangles of the same size were used. Pixel intensity of *Rnd3* stain was measured and expressed as the ratio sulcus/gyrus signal for each region.

### RNAseq analysis

Single cell RNAseq data for mouse cortex samples were obtained from the published NCBI Gene Expression Omnibus with accession numbers GSE65000^63^ and GSE153164^64^. Data from GSE6500 was retrieved directly from NCBI and the expression level of the mouse *Rnd3* (ENSMUSG00000017144) or *FLRT3* (ENSMUSG00000051379) were normalized by *GAPDH* (ENSMUSG00000057666) for each data set: 4 replicates for aRG (apical progenitors, AP) and 4 replicates for neurons (N). Data for GSE153164 was retrieved from the Single Cell Portal (Broad Institute https://singlecell.broadinstitute.org/single_cell/study/SCP1290/molecular-logic-of-cellular-diversification-in-the-mammalian-cerebral-cortex) using the UMAP clustering and filtering for general cell type (apical and basal progenitors, interneurons, migrating neurons and projecting neurons) and E15 mouse data.

### Statistical analysis

One-tail unpaired Student’s *t* test, one-way ANOVA or two-way ANOVA with Tukey post-hoc analysis (GraphPad Prism 8.0), were used to determine statistical differences. Significance was considered when p <0.05 (*), p <0.01 (**) or p <0.001 (***). Error bars were calculated using the standard error of the mean (SEM).

## Results

### *Rnd3* expression in the developing cortex

To investigate the function of Rnd3 during cortex formation we first studied its expression at different developmental stages. Unfortunately, none of the commercially available antibodies that we tried showed specific signal when compared to knock-out tissue (data not shown). Instead, we took advantage of a gene trap (gt) allele of *Rnd3* that contains the bacterial *lacZ* gene and expresses β-galactosidase (β-GAL) under the endogenous *Rnd3* promoter (see Methods) and which reliably reproduces *Rnd3* gene expression^51^. We therefore obtained sections from heterozygous *Rnd3^gt/+^* brains at embryonic (E) days 12.5, 14.5, 16.5 and 18.5, which were stained with X-Gal. We observed that *Rnd3* expression in the cortex was very dynamic. At E12.5, βGAL activity was very low but increased greatly at E14.5 and E16.5 while at E18.5 became fainter (Figure 1a-d). We immunostained these sections with neuronal markers such as CTIP2 or TBR1 or progenitor cells such as TBR2 (BPs) or SOX2 (apical progenitors). At E13.5 and E14.5, βGAL+ cells were observed exclusively in the intermediate zone (IZ), right below the cortical plate (CP) which co-expresses TBR1 and CTIP2 at these stages (Figure 1e-h’’’). At E16.5, *Rnd3* expression became broader and βGAL+ cells were detected, in addition to the IZ, in the SVZ and in the VZ, but not in the CP (Figure 1i,j’’’). At E18.5, βGAL+ signal was only present in progenitor cells in the SVZ (TBR2+;SOX2-) and in the VZ (TBR2-;SOX2+) (Figure 1k,l’’’). These results show that RND3 is mainly expressed in migrating neurons within the IZ at early stages (E13.5-E14.5), in migrating neurons and progenitor cells at E16.5 and only in progenitor cells at E18.5. Expression of *Rnd3* in migrating neurons was confirmed by showing co-expression at E14.5 in *Rnd3^gt/+^* brain sections of βGAL and CTIP2 within the IZ, as migrating neurons in this region also express CTIP2, albeit at lower levels than neurons in the CP^65^ (Figure 1m-m’’). Taken together, these results suggest a role of RND3 in cortical development, mainly in early neuron migration (E13.5-E16.5) and/or late proliferation (E16.5-E18.5).

**Figure 1:**
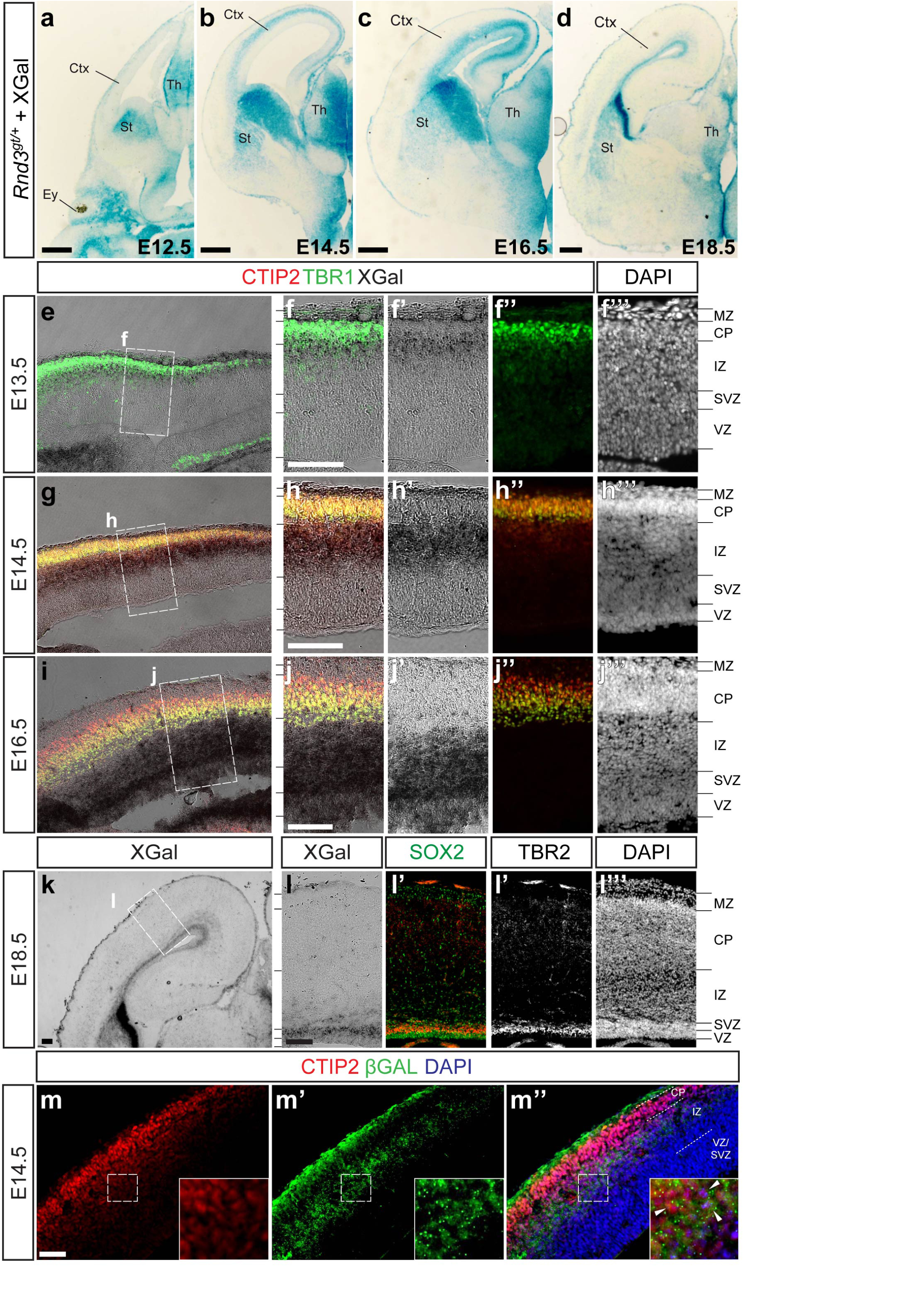
Expression of *Rnd3* during mouse cortex development. **a-d** XGal staining (*Rnd3* expression) of coronal sections of *Rnd3^gt/+^*brains at different developmental stages (E12.5-E18.5, as indicated). Only one hemisphere is shown. **e** XGal staining (*Rnd3* expression) and TBR1 staining of a coronal section of the cortex of a *Rnd3^gt/+^*brain, at E13.5. Dashed box (**f**) indicates the position of panels **f-f’’’**. **f**-**f’’’** High magnification of the area depicted in **e**, showing TBR1+XGal (F), XGal (**f’**), TBR1 (**f’’**) and DAPI (**f’’’**) staining. **g** XGal staining (*Rnd3* expression) and TBR1+CTIP2 double staining of a coronal section of the cortex of a *Rnd3^gt/+^* brain, at E14.5. Dashed box (**h**) indicates the position of panels **h**-**h’’’**. **h**-**h’’’** High magnification of the area depicted in **g**, showing TBR1+CTIP2+XGal (**h**), XGal (**h’**), TBR1+CTIP2 (**h’’**) and DAPI (**h’’’**) staining. **i** XGal staining (*Rnd3* expression) and TBR1+CTIP2 double staining of a coronal section of the cortex of a *Rnd3^gt/+^* brain, at E16.5. Dashed box (**j**) indicates the position of panels **j**-**j’’’**. **j**-**j’’’** High magnification of the area depicted in **i**, showing TBR1+CTIP2+XGal (**j**), XGal (**j’**), TBR1/CTIP2 (**j’’**) and DAPI (**j’’’**) staining. **k** XGal staining (*Rnd3* expression) of a coronal section of the cortex of a *Rnd3^gt/+^* brain, at E18.5. Dashed box (**k**) indicates the position of panels **l**-**l’’’**. **l**-**l’’’** High magnification of the area depicted in **k**, showing XGal (**l**), SOX2 (**l’**), TBR2 (**l’’**) and DAPI (**l’’’**) staining. **m**-**m’’** CTIP2 (**m**) and βGAL (**m’**) double staining (merged in **m’’**) of a coronal section of the cortex of a *Rnd3^gt/+^*brain, at E14.5. Insets at the bottom-right are magnified images of the area depicted by dashed boxes in each panel. Arrowheads in **m’’** indicate the presence of βGAL (*Rnd3* expression) in low CTIP2-expressing cells in the IZ. The MZ (marginal zone), CP (cortical plate), IZ (intermediate zone), SVZ (subventricular zone) and VZ (ventricular zone) are indicated in panels **f’’’**, **h’’’**, **j’’’**, **l’’’**. Scale bars: **a**-**d**, 400 μm; **f**,**h**,**j**,**k**,**l**,**m**, 200 μm.

### Genetic deletion of *Rnd3* results in the appearance of spontaneous cortical sulci

To address the function of RND3 during cortex development in vivo, we generated homozygous *Rnd3^gt/gt^* embryos to study their cortical structure at different embryonic stages. Surprisingly, spontaneous cortical sulci appeared in some of the mutant embryos: in 1 out of 12 at E14.5, in 2 out of 9 at E16.5 (Figure 2a,b,h) and in 3 out of 14 at E18.5 (Figure 2c,d,h). Interestingly, while none of the *Rnd3^+/+^* littermates presented cortical sulci (0 out of 20, Figure 2h), some of the heterozygous *Rnd3^gt/+^* embryos did (2 out of 27, Figure 2h), although the penetrance was lower than in *Rnd3^gt/gt^* (6 out of 35, Figure 2h). This observation is suggestive of a gene dose effect of *Rnd3* during cortex development. Like in gyrencephalic species, the observed sulci involved mainly the pial but not the apical surface (arrowheads in Figure 2)^66^.

**Figure 2:**
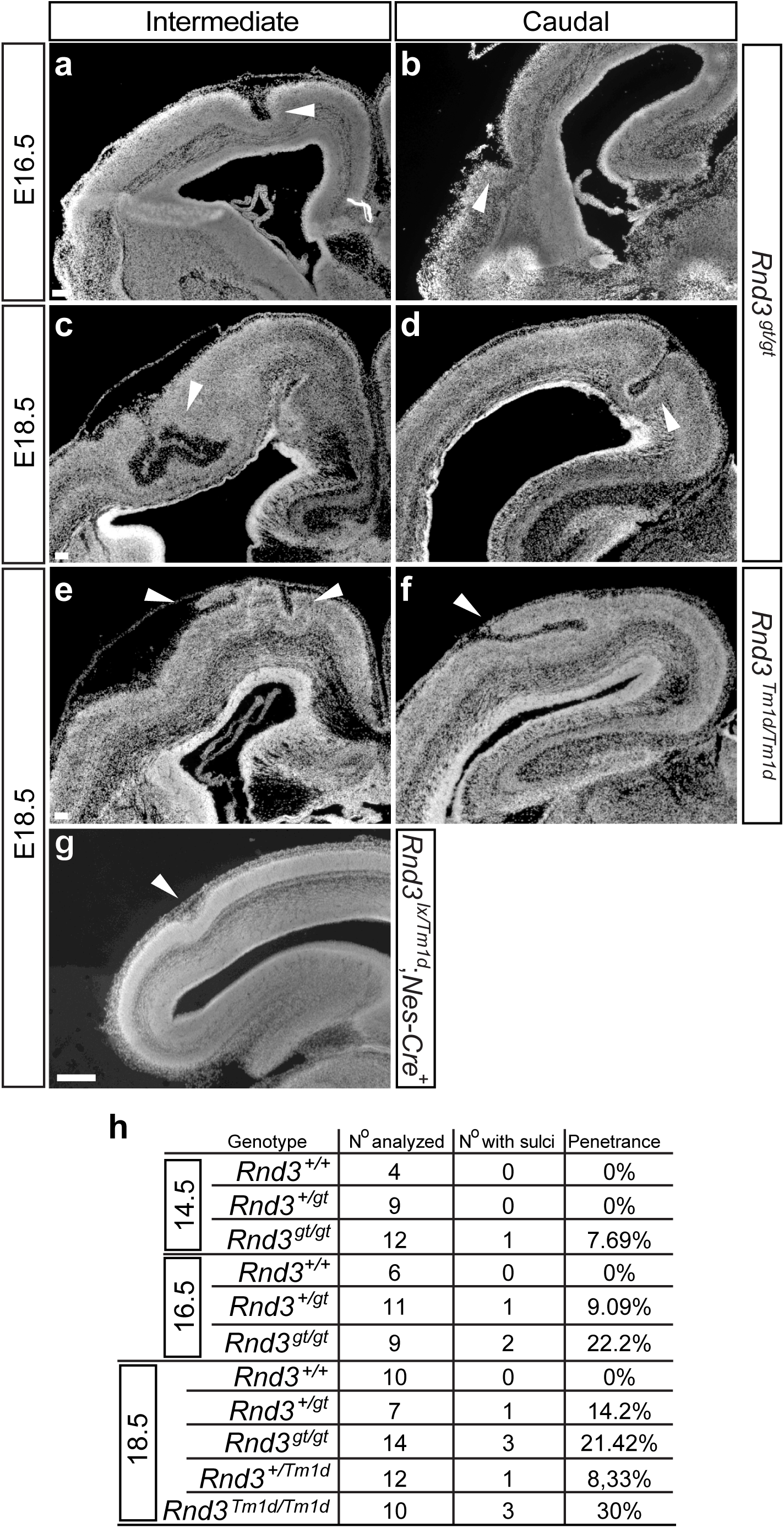
Abnormal development in *Rnd3* knock-out brains reveals the formation of spontaneous cortical sulci. **a-g** DAPI staining of cortical sulci (arrowheads) of coronal sections of the cortex of *Rnd3^gt/gt^*(**a**-**d**), *Rnd3^Tm1d/Tm1d^* (**e**,**f**), *Rnd3^lx/Tm1d^*;*Nestin-Cre*^+^ (Nes-Cre, **g**) brains at intermediate (**a**,**c**,**e**,**g**) and caudal (**b**,**d**,**f**) positions and at different developmental stages (**a**,**b**: E16.5; **e**-**g**: E18.5). **h** Summary table of the number and percentage of *Rnd3* mutant embryos displaying cortical folds for the indicated genotypes, at E14.5, E16.5 and E18.5. Scale bar, 200 μm.

We also generated an independent *Rnd3* null allele (*Rnd3^Tm1d^*) using a deleter Cre line and a conditional, *Rnd3* floxed allele (*Rnd3^lx^*) (see Methods and reference 51). As expected, E18.5 *Rnd3^Tm1d/Tm1d^* embryos also displayed cortical folds with partial penetrance (3 out of 10, Figure 2e,f,h). Moreover, we crossed homozygous *Rnd3^lx/lx^* females with heterozygous *Rnd3^Tm1d/+^* males harbouring a nervous system specific Cre (Nestin-Cre^67^) to obtain *Rnd3^lx/Tm1d^;Nestin-Cre^+^* embryos with specific deletion of *Rnd3* in the nervous system. Interestingly, 3 out 15 of these mutant embryos (20%) also displayed cortical folds indicating that this defect is largely nervous system specific (Figure 2g). Since the cortical folds that appeared in *Rnd3^gt/gt^* or *Rnd3^Tm1d/Tm1d^*mutants were indistinguishable, both type of embryos was pooled for the rest of the analysis and were considered as *Rnd3^KO^*. Finally, we analysed the preferred region for cortical folding and noticed that all of the cortical folds were observed in intermediate and/or caudal positions of the mutant brains: 25% (3/12) were found in the dorsal part (as in Figure 2d), 58,3% (7/12) in the lateral part (as in Figure 2b) and 16,7% (2/12) in both, dorsal and lateral (as in Figure 2e,f). In summary, these results show that *Rnd3* mutants display spontaneous cortical sulci in intermediate/caudal parts of the cortex, a defect that is nervous-system specific.

### Proliferation is not affected and cortical structure is largely preserved in the *Rnd3^KO^* brains

To start investigating the mechanisms underlying the formation of cortical folds in the *Rnd3* mutants, we first analysed if cell proliferation was increased, as an expansion of BPs during development is one of the mechanisms known to promote cortical folding^8,9^. Moreover, RND3 has been reported to inhibit BPs proliferation^45^. From our expression analysis based on X-Gal staining (Figure 1), migrating neurons show higher *Rnd3* expression at earlier stages (E13.5-E14.5), whereas at later time points the signal appears to be mainly restricted to progenitors (E16.5-E18.5). We therefore studied proliferation effects in the *Rnd3^KO^* cortices at late embryonic stages. For that we used the M phase mitotic marker phospho-histone H3 (PH3^68^) and stained control and *Rnd3^KO^* brain sections with folds, at E18.5 (Figure 3a-d). We then quantified the total number of PH3+ cells per brain (Figure 3e), the number of PH3+ cells per section (distinguishing those containing folds from those without folds, in the *Rnd3^KO^* brains, Figure 3f) and the relative distribution of PH3+ cells in the VZ, SVZ, IZ and CP along the rostro-caudal axis of the brain (Figure 3g). Noticeably, we did not detect a significant increase in any of the quantifications that we made, arguing against a major role of RND3 in inhibiting cell proliferation that could contribute to the formation of the cortical folds in the *Rnd3^KO^* brains.

**Figure 3:**
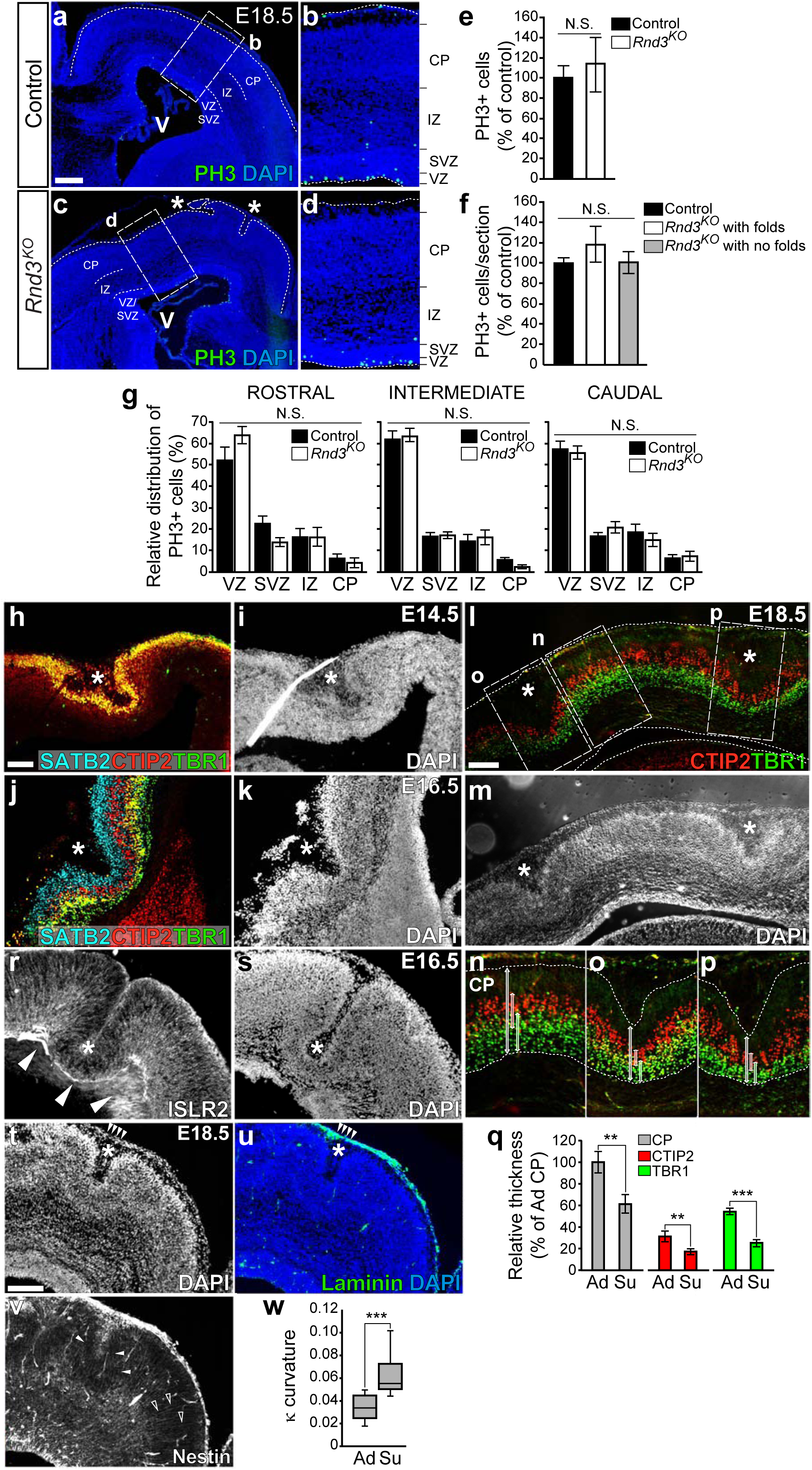
Proliferation is not affected in *Rnd3^KO^* cortices and cortical structure is largely preserved. **a-d** PH3 and DAPI staining of coronal sections of the cortex of control (**a**,**b**) and *Rnd3^KO^* brains (**c**,**d**), at E18.5. Cortical sulci in the *Rnd3^KO^*brain are indicated with asterisks (**c**). Dashed squares in **a** and **c** show the position of the high magnification images in **b** and **d**, respectively. CP, cortical plate; MZ, marginal zone; IZ, intermediate zone; VZ/SVZ, ventricular and subventricular zone; V, ventricle. Scale bar, 200 μm. **e** Quantification of the total PH3+ cells along the rostro-caudal axis of control (black bar) and *Rnd3^KO^* cortices (white bar), at E18.5. Values are expressed as percentage respect to controls (100%) and as the mean±SEM; n=3 brains/genotype, >10 sections/brain. No statistical differences (N.S., one-tail Student’s *t* test, p=0.351). **f** Quantification of the PH3+ cells/section in cortices of controls (black bar), *Rnd3^KO^* with folds (white bar) and *Rnd3^KO^* without folds (grey bar), at E18.5. Values are expressed as percentage respect to controls (100%) and as the mean±SEM; n>9 sections from 3 brains/genotype. No statistical differences (N.S., one-way ANOVA, p=0.459). **g** Quantification of the relative distribution (%) of PH3+ cells in the ventricular zone (VZ), subventricular zone (SVZ), intermediate zone (IZ) and cortical plate (CP) within rostral, intermediate and caudal sections of control (black bars) and *Rnd3^KO^*(white bars) cortices, at E18.5. Values are expressed as the mean±SEM; n>9 sections from 3 brains/genotype. No statistical differences d (N.S., two-way ANOVA, p=0.719, intermediate; p=0.553, caudal). N.S.*, statistical differences by two-way ANOVA in rostral sections (p=0.02), but not after Tukey post-hoc analysis between genotypes. **h**-**k** SATB2 (turquoise), CTIP2 (red) and TBR1 (green) (**h**,**j**) or DAPI (**i**,**k**) staining of cortical folds (asterisks) of coronal sections of *Rnd3^KO^* brains at E14.5 (**h**,**i**) oe E16.5 (**j**,**k**). **l**,**m** CTIP2 (red) and TBR1 (green) (**l**) or DAPI (**m**) staining of two simultaneous cortical sulci (asterisks) on a coronal section of a *Rnd3^KO^* brain, at E18.5. Dashed boxes in **l** indicate the position of panels **n**-**p**. **n**-**p** High magnification of the two folds depicted in **l** (**o**,**p**) and of the adjacent region with no folds (**n**), stained with CTIP2 (red) and TBR1 (green). The two dotted lines in each panel defines the boundaries of the CP. Bidirectional arrows in grey, red and green indicate the width of the CP, CTIP2 and TBR1 regions, respectively. **q** Quantification of the width of the CP (grey bars), CTIP2 (red bars) and TBR1 (green bars) staining in sulci (Su) or in adjacent region (Ad), as in **n**-**p**. Values are expressed as percentage respect to the width of the CP in the adjacent region (100%) and as the mean±SEM; n>3 for each region. **<p=0.01; ***<p=0.001, one-tail Student’s *t* test. **r**,**s** ISLR2 (**r**) or DAPI (**s**) staining of cortical folds on coronal sections of *Rnd3^KO^* brains at E16.5. **t**-**v** DAPI (**t**), laminin (green) and DAPI (blue) (**u**) and nestin (**v**) staining of a cortical sulcus (asterisk) on a coronal section of *Rnd3^KO^* brain at E18.5. Arrowheads in **t**,**u** indicate the superficial laminin staining in the sulcus. Arrowheads and empty arrowheads in **v** indicate the curved and the straight trajectories of the radial glial processes around the sulcus and in an adjacent area without folding, respectively. **w** Box-and-whisker plots showing the quantification of the curvature of the radial glial processes stained with nestin (as in **v**) in sulcus (Su) or adjacent region (Ad), as indicated, using the kappa plugin (κ) in ImageJ. n>12 processes from three sulci. ***p<0.001, one-tail Student’s *t* test. Scale bar, 200 μm.

We next characterized the structure of the cortex of *Rnd3^KO^*brains. We realized that overall cortical width was slightly thinner in *Rnd3^KO^* brains with folds than in *Rnd3^KO^* brains without folds or controls at E16.5 and E18.5, especially in rostral and intermediate regions (Supplementary Figure 1a-d and data not shown). However, this was not the result of a massive reduction of cells as the total cell count, by DAPI staining, was similar among the three groups of embryos, except in rostral regions where a slightly decrease of DAPI counts was observed in *Rnd3^KO^* with folds compared to controls (p=0,03143, One-way ANOVA with Tukey post-hoc analysis, Supplementary Figure 1e).

Then we studied the layering organization within the CP of the *Rnd3^KO^* cortex by immunolabeling against SATB2 (upper CP), CTIP2 (middle CP) or TBR1 (lower CP) and observed that their relative position was largely preserved (Supplementary Figure 1f-k’’’). Layering organization was also intact within the folds at E14.5, E16.5 and E18.5 (Figure 3h-m), as observed in gyrencephalic brains^8^. Interestingly, there was a significant reduction of the width of CTIP2 and TBR1 layers within the sulcus compared to adjacent areas, similar to what was reported in gyrencephalic species such as the ferret^69^ (Figure 3n-q). In addition, the corticofugal axons, immunostained against the transmembrane protein ISLR2^51^ or against neurofilament, were correctly emerging from the sulci (Figure 3r,s and data not shown). We also immunostained these sulci with nestin and laminin at E18.5 (Figure 3t-v) to study the morphology of the radial glial processes and the meningeal basement membrane around the folds, respectively. We observed that the curvature index (κ) of the radial glial processes was significantly higher in the sulcus than in adjacent areas due of the convergence of fibers at sulcal pits (Figure 3v,w), like those reported in gyrencephalic species such as ferrets and monkeys^69,70^. The meningeal basement membrane, stained with laminin was smooth and intact indicating that the sulci were not the result of neuronal ectopias (Figure 3t,u). Altogether, these results indicate that cortical structure is largely preserved in the *Rnd3^KO^* brains and that the folds display most of the features of the gyrencephalic folds therefore suggesting that RND3 might be an important regulator of cortical folding in vivo.

### Cortical radial migration is regulated by RND3

Remarkably, *Rnd3* is expressed in the IZ from E14.5 until E16.5 (Figure 1), suggesting a role of RND3 during cortical neuron migration. In fact, RND3 has been previously shown to regulate locomotion within the CP during cortex development^44^. Thus, we addressed the involvement of RND3 in cortical radial migration by altering RND3 expression levels by IUE. Embryos were IUE at E13.5 and analysed at E19.5 by immunofluorescence on brain sections against the reporter gene EGFP to visualize the localization of the electroporated cells within the cortex (see Methods). Three experimental conditions were considered: control embryos, electroporated with the empty vector (only expressing EGFP), overexpression of RND3 (gain-of-function condition) or electroporation of CRE on *Rnd3^lx/lx^* embryos to ablate *Rnd3* expression (loss-of-function condition).

To facilitate the quantification of the distribution of the EGFP+ cells, the cortex width was divided into 10 equal bins. In control embryos, most of the EGFP+ cells were observed in upper parts of the cortex, as expected (Figure 4a,d). In contrast, overexpression of RND3 caused a significant shift towards deeper regions (Figure 4b,d). Consistently, ablation of *Rnd3* induced the opposite effect and more EGFP+ cells were observed in upper parts of the cortex (Figure 4c,d). Importantly, the total number of EGFP+ cells among the three conditions was similar and no significant differences were observed (Figure 4e) arguing against a role of RND3 on cell proliferation and/or viability. Instead, these results indicate that RND3 modulates the radial migration of cortical neurons during development.

**Figure 4:**
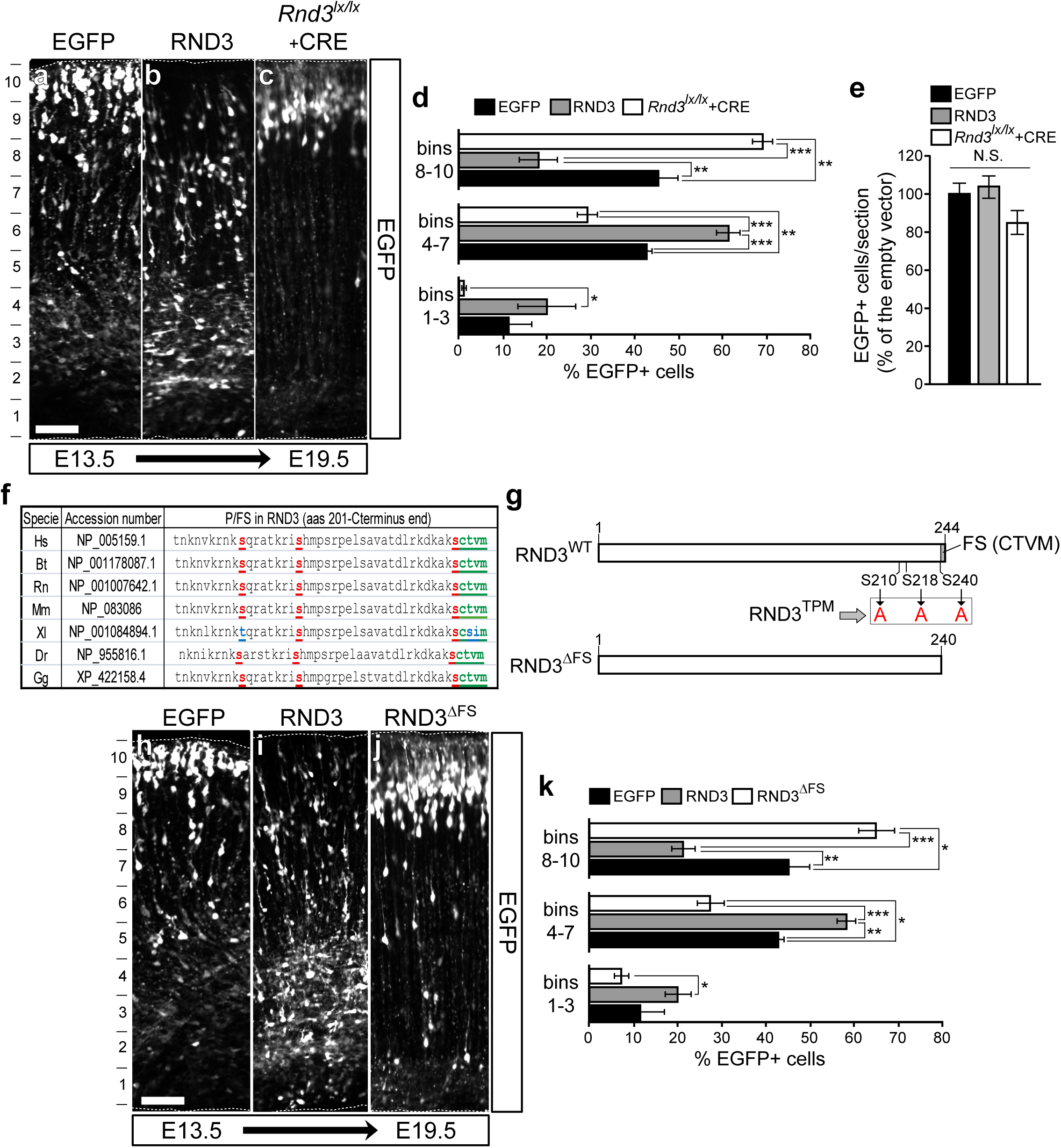
Rnd3 regulates cortical neuron migration during development and requires an intact farnesylation motif. **a-c** EGFP staining of coronal sections of the cortex of wild-type (**a**,**b**) and *Rnd3^lx/lx^* brains (**c**) IUE with empty vector (**a**, EGFP), RND3 (**b**) or CRE (**c**) at E13.5 and analysed at E19.5. The distribution of the electroporated cells was quantified through the cortical wall divided in ten equal bins (left). Scale bar, 100 μm. **d** Quantification of the relative distribution (%) of EGFP+ cells in lower (1-3), middle (4-7) and upper (8-10) cortical bins of wild-type embryos IUE with empty vector (EGFP, black bars) or with RND3 (grey bars) or of *Rnd3^lx/lx^* embryos IUE with CRE (white bars). Values are expressed as the mean±SEM; n>3 brains/condition, >5 sections/brain. *<p=0.05; **<p=0.01; ***<p=0.001, one-way ANOVA with Tukey post-hoc analysis. **e** Quantification of the EGFP+ cells/section in cortices of wild-type embryos IUE with empty vector (EGFP, black bars) or RND3 (grey bars) or of *Rnd3^lx/lx^* embryos IUE with CRE (white bars). Values are expresses as percentage respect to controls (100%) and as the mean±SEM; n>3 brains/condition, >5 sections/brain. No statistical differences (N.S., one-way ANOVA, p=0.0724). **f** Conservation of the phosphorylation (P) sites S210, S218 and S240 (in red) and the farnesylation site (FS, in green) at C-terminus of RND3 among different vertebrate species. Bt, *Bos taurus*; Gg, *Gallus gallus*; Dr, *Danio rerio*; Hs, *Homo sapiens*; Mm, *Mus musculus*; Rn, *Rat norvegicus*; Xl, *Xenopus laevis*. In Xl, the conserved threonine in the FS is substituted by serine (S in blue). **g** Scheme of the RND3 protein with the position of the phosphorylation sites S210, S218 and S240 and the CAAX motif (CTVM) and showing the mutants generated in this study: the RND3^TPM^, triple phosphorylation mutant in which S210, S218 and S240 were substituted by the non-phosphorylatable amino acid alanine (A) and the RND3^ΔFS^, in which a STOP codon was introduced right before the last four aminoacids containing the farnesylation motif (FS). **h**-**j** EGFP staining of coronal sections of the cortex of wild-type brains IUE with empty vector (**h**, EGFP), RND3 (**i**) or RND3^ΔFS^ (**j)** at E13.5 and analyzed at E19.5. The distribution of the electroporated cells was quantified through the cortical wall divided in ten equal bins (left). Scale bar, 100 μm. **k** Quantification of the relative distribution (%) of EGFP+ cells in lower (1-3), middle (4-7) and upper (8-10) cortical bins of wild-type embryos IUE with empty vector (EGFP, black bars), RND3 (grey bars) or RND3^ΔFS^ (white bars). Values are expressed as the mean±SEM; n>3 brains/condition, >5 sections/brain. *<p=0.05; **<p=0.01; ***<p=0.001, one-way ANOVA with Tukey post-hoc analysis.

We next investigated the signalling mechanisms involved in the control of neuron migration by RND3. Since RNDs are constitutive active proteins, they have been shown to be regulated by posttranslational modifications such as farnesylation and phosphorylation^34^. RND3 contains a conserved farnesylation motif at the C-terminus end with a cysteine, two aliphatic aminoacids and any aminoacid (CAAX motif) and three conserved phosphorylation sites, at serines 210, 218 and 240 (Figure 4f)^33,35,71,72^. To address the relevance of these post-translational modifications for RND3 function during neuron migration, we generated RND3 mutants in which the farnesylation motif was deleted by adding a premature stop codon (RND3^ΔFS^) or in which the three phosphorylation sites were mutated to alanine (RND3^TPM^; Figure 4g and Methods). These constructs were well expressed and often localized nearby the plasma membrane in heterologous cells, except for RND3^ΔFS^ which, as expected, lost its membrane tethering and showed a more diffused intracellular pattern (Supplementary Figure 2a-f). Then, we IUE these constructs in wild-type embryonic cortical brains and observed that RND3^ΔFS^ abolished the effects on neuron migration of RND3, suggesting that membrane localization is essential for RND3 function (Figure 4h-k). On the other hand, IUE of RND3^TPM^ also reduced migration effects of RND3, although at lesser extent (Supplementary Figure 2g-j). Taken together these results indicate that RND3 regulates neuron migration during cortex development and that membrane localization and phosphorylation at serines 210, 218 and 240, are required for this function.

### RND3 interacts with FLRT3 and both genes are co-expressed in migrating cortical neurons during development

The fine tuning of RND function at the plasma membrane has been shown to be modulated by the interaction with specific transmembrane receptors such as PlexinBs^52,53^ or Notch^41^. On the other hand, FLRT3, was shown to interact with RND1 downstream of TGF-β signalling and modulate cell adhesion during gastrulation in Xenopus embryos^55^. However, the relevance of the FLRT-RND interaction in the nervous system and cortex development is currently unknown. Interestingly, *Flrt1/3* DKOs also display spontaneous cortical sulci with intriguingly similar features to those observed here in the *Rnd3^KO^* embryos^26^. These observations prompted us to investigate a functional relationship between FLRTs and RND3 during cortex development. To start exploring this possibility, we first tested the interaction of RND3 with each of the FLRT proteins by co-immunoprecipitation in transfected HEK293T cells and observed a stronger interaction with FLRT3 than with FLRT1 or FLRT2 (Supplementary Figure 2k and data not shown). As expected, this interaction required an intact FLRT3 intracellular domain (ICD), as its substitution by EGFP (FLRT3^ΔCEGFP^)^25^, abolished its co-immunoprecipitation with RND3 (Supplementary Figure 2k). We then focused on this FLRT3-Rnd3 interaction and observed that both genes were strongly expressed in the IZ during cortex development, an area enriched of migrating neurons (Figure 5a,b). Using available data from two single-cell RNA sequencing (sc-RNAseq) databases we confirmed that *Flrt3* and *Rnd3* were both indeed expressed in migrating neurons (Figure 5c-f)^63,64^. In the *Flrt1*/*3* DKOs, cortical folding is caused by a decrease of FLRT-mediated homophylic cell adhesion triggering a reduced intercellular adhesion with increase neuron migration and clustering in the cortical plate^26^. However, it is not known if the cytoplasmic domain of FLRTs is necessary for this effect and, if so, which are the downstream intracellular mechanisms involved. Interestingly, IUE of FLRT3 caused a delay of neuron migration like RND3 overexpression while the IUE of FLRT3^ΔCEGFP^ almost completely abolished this effect, like *Rnd3* ablation (Figure 5g-j). These results suggest that the control of cortical neuron migration by FLRT3 requires an instructive signalling activity triggered by an intact ICD. Considering the interaction of RND3-FLRT3, their co-expression in migrating cortical neurons, their similar effects on neuron migration by IUE and the comparable cortical folding phenotype in knock-out embryos, we hypothesized that FLRT3 and RND3 are part of the same signalling pathway during cortex development regulating the formation of cortical folds.

**Figure 5:**
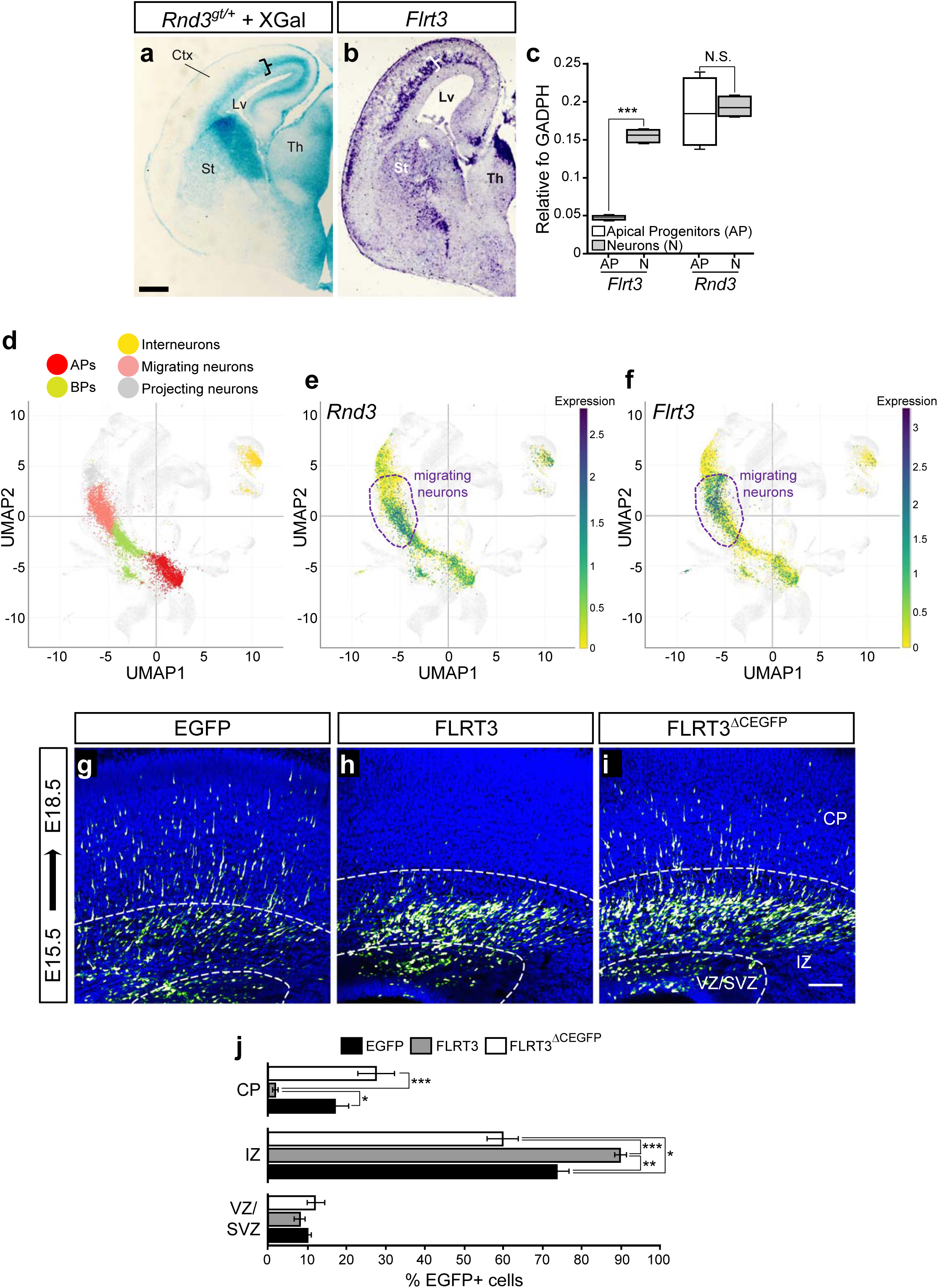
*Rnd3* and *FLRT3* are co-expressed in migrating cortical neurons. **a,b** XGal staining (*Rnd3* expression) (**a**) and in situ hybridization of *Flrt3* (**b**) of coronal sections of *Rnd3^gt/+^*or wild-type brains, respectively, at E16.5. Ctx, cortex; Lv, lateral ventricle; St, striatum; Th, thalamus. Braces in the cortex indicate the highest expression of *Rnd3* and *FLRT3*, both in the IZ. Scale bar, 400 μm. **c** Box-and-whisker plots showing *Flrt3* and *Rnd3* mRNA expression levels normalized to *GADPH* mRNA in neurons (N) or apical progenitors (APs), using published mRNA profiling from E14.5 mouse embryos (GSE65000^63^). *Flrt3* and *Rnd3* mRNAs are both abundant in neurons (N). *Flrt3* mRNA is less expressed in APs (n=4 replicates; ***p<0.001, one-tail Student’s *t* test) while *Rnd3* mRNA is similarly expressed in APs and neurons (n=4 replicates; N.S., no statistical difference, p=0,392). **d** UMAP visualization of the published single cell *RNA* data (Single Cell Portal, Broad Institute) from E15.5 mouse cortex. Five major cell clusters are coloured as indicated by cell-type assignment based on published metadata (GSE153164^64^) (BP, basal progenitors; AP, apical progenitors). **e**,**f** UMAP plots of *Flrt3* (E) and *Rnd3* (F) mRNA expression per cell (GSE153164^64^). Expression level is illustrated on the right. Most of the *Flrt3*-expressing cells belong to the migrating neuron cluster (dashed violet line) while *Rnd3*-expressing cells are mainly distributed between migrating neurons (dashed violet line) and APs. **g**-**i** EGFP (green) and DAPI (blue) staining of coronal sections of the cortex of wild-type bains IUE with empty vector (**g**, EGFP), FLRT3 (**h**) or FLRT3 mutant in which the intracellular domain was substituted by EGFP (FLRT3^ΔCEGFP^, **i**) at E15.5 and analysed at E18.5. The distribution of the electroporated cells was quantified through the cortical wall divided in VZ/SVZ, IZ and the CP. Scale bar, 100 μm. **j** Quantification of the relative distribution (%)of EGFP+ cells in VZ/SVZ, IZ and CP of wild-type embryos IUE with empty vector (EGFP, black bars), FLRT3 (grey bars) or FLRT3^ΔCEGFP^ (white bars). Values are expressed as the mean±SEM; n>4 brains/condition, >3 sections/brain. *<p=0.05; **<p=0.01; ***<p=0.001, one-way ANOVA with Tukey post-hoc analysis.

### Rnd3 interaction with FLRT3 is necessary for proper cortex development

In order to investigate the possibility that RND3 and FLRT3 could be part of the same signalling pathway regulating cortex development and folding we first modelled the interaction of the ICD of FLRT1-3 and RND1-3 *in sillico* using the artificial intelligence-based platform AlphaFold 3.0. The aim was to predict the interacting surfaces and generate structure-based mutants to impair binding during cortical development. In the resulting models, all predicted complexes of RND3 and FLRTs involved a stretch of amino acids lining the central β-sheet (sequence FENYTASF, Figure 6a,b and Supplementary Figure 3a,b). This sequence also provides most interactions in the models calculated for RND1 and FLRT1-3 (Supplementary Figure 3c). The sequence is conserved across different animal species (Supplementary Figure 3d) and overlaps with a known binding motif that engages plexin receptors (PDB code 2REX)^73^. We attempted to calculate models also for RND2 and FLRTs (Supplementary Figure 3c) but note that this resulted in twisting of one of the beta-strands within the core domain of RND2, which is likely an artefact of the method. Conversely to RND3, where the same interaction surface was predicted for all three complexes, the corresponding interaction motif on FLRTs was not the same in each complex (Supplementary Figure 4a,b). This is in agreement with the overall low level of sequence conservation for FLRT intracellular domains. We focused on RND3, which displays the conserved binding motif to FLRTs encompassing the amino acids FENYTASF and mutated this motif to **<u>PGTP</u>**T**<u>PPP</u>**with the aim to generate an RND3 mutant (RND3^F3BM^) unable to bind to FLRT3^ICD^ (Figure 6c). Calculated models suggest that, apart from local changes where the mutations were introduced, the overall fold of the protein is not disrupted in this mutant (Supplementary Figure 4c,d). RND3^F3BM^ was well expressed in HEK293T cells and was properly localized within the cell as the wild-type RND3, including the plasma membrane, although it displayed a slightly higher apparent molecular weight in a Western blot when compared to the wild-type protein (Supplementary Figure 2a). As expected, when RND3^F3BM^ was co-expressed with FLRT3, it failed to bring down FLRT3 in a co-immunoprecipitation assay (Figure 6d) which validated this binding mutant. Next, we used IUE to assess the effects of this RND3^F3BM^ during cortical neuron migration and, strikingly, we found that it failed to induce the delay effect triggered by RND3 (Figure 6e-h). This is a very interesting observation as it strongly suggests that RND3 is an intracellular downstream effector of FLRT3 for cortical neuron migration during cortex development. Surprisingly, we also noticed that most of the embryos IUE with the RND3^F3BM^ (6 out of 7), displayed cortical sulci in the electroporated area, indicating that FLRT3-RND3 interaction is important for the control of cortical folding (Figure 6i-l).

**Figure 6:**
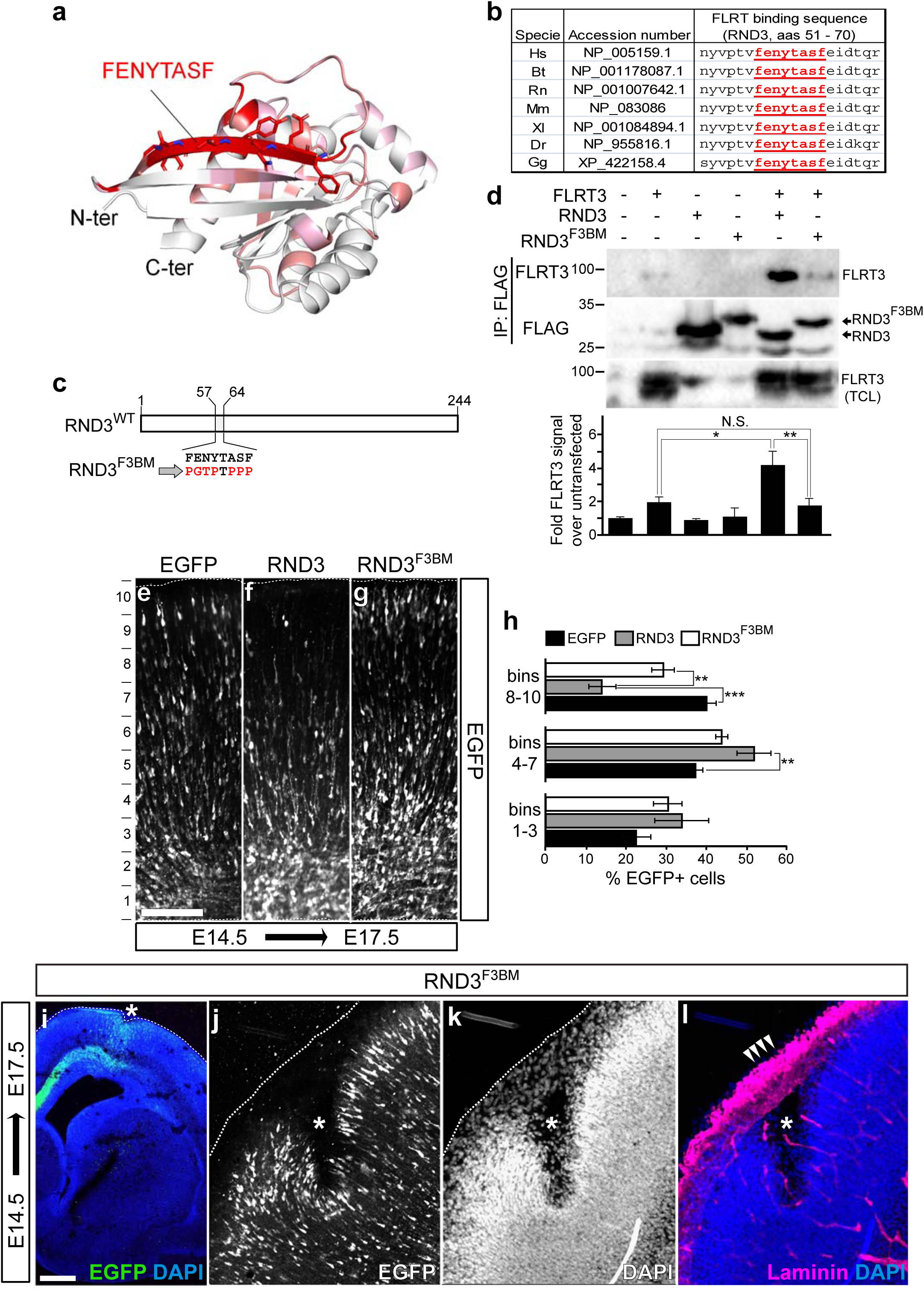
The interaction of Rnd3 and FLRT3 is important for neuron migration during cortex development. **a** Predicted model of RND3 using AlphaFold 2.0. The binding surface to FLRTs is shown in red and encompasses the amino acid sequence FENYTASF. **b** Summary table of the protein sequence between aminoacids (aas) 51 and 70 of RND3 of different vertebrate species encompassing the conserved FLRT binding sequence (FENYTASF in red), predicted by AlphaFold 2.0. Bt, *Bos taurus*; Gg, *Gallus gallus*; Dr, *Danio rerio*; Hs, *Homo sapiens*; Mm, *Mus musculus*; Rn, *Rat norvegicus*; Xl, *Xenopus laevis*. **c** Scheme depicting the position of the RND3 binding sequence to FLRT3 and the mutant generated in this study to prevent RND3-FLRT3 interaction (Rnd3^F3BM^), in which the FENYTASF sequence was mutated to PGTPTPPP (in red). **d** Co-immunoprecipitation assay in transfected HEK293T cells with the indicated constructs. The lysates were immunoprecipitated with FLAG antibodies to pull down RND3 proteins and analyzed by Western-blot with anti-FLRT3 (upper panel) or anti-FLAG antibodies (middle panel). The input levels of FLRT3 in the total cell lysates (TCL) are shown (lower panel). Molecular weight markers are indicated on the left. The bottom graph shows the quantification of the co-immunoprecipitated FLRT3. Values are expressed as fold increase respect to the non-transfected cells (1), normalized by the total levels of RND3 (immunoprecipitates) and FLRT3 (TCL), and as the mean±SEM; n=4 experiments. *p<0.05, **p<0.01, one-way ANOVA with Tukey post-hoc analysis; N.S., no statistical difference, p=0.999. **e**-**g**) EGFP staining of coronal sections of the cortex of wild-type brains IUE with empty vector (**e**, EGFP), RND3 (**f**) or RND3^F3BM^ (**g**) at E14.5 and analysed at E17.5. The distribution of the electroporated cells was quantified through the cortical wall divided in ten equal bins (left). Scale bar, 100 μm. **h** Quantification of the relative distribution (%) of EGFP+ cells in the lower (1-3), middle (4-7) and upper (8-10) cortical bins of wild-type embryos IUE with empty vector (EGFP, black bars), RND3 (grey bars) or RND3^F3BM^ (white bars). Values are expressed as the mean±SEM; n>3 brains/condition, >5 sections/brain. *<p=0.05; **<p=0.01; ***<p=0.001, one-way ANOVA with Tukey post-hoc analysis. **i**-**l** EGFP (green) and DAPI (blue) (i), EGFP (j), DAPI (k), laminin (magenta) and DAPI (blue) staining of cortical sulcus (asterisk) on a coronal section of wild-type brain IUE with RND3^F3BM^ at E14.5 and analysed at E17.5. Arrowheads in **l** indicate the superficial laminin staining in the sulcus. Scale bar, 200 μm.

### *Rnd3* is downregulated in the OSVZ of prospective sulci in a naturally gyrencephalic animal model

Previous results show a negative correlation of *Flrt1* and *Flrt3* expression levels with the formation of cortical folds in gyrencephalic species. In ferret, for instance, *Flrt1* and *Flrt3* mRNAs are less abundant in prospective sulcus than in gyrus areas at the onset of folding development^26^. This agrees with the fact that ablation of *Flrt1*/*Flrt3* in mice allows the manifestation of spontaneous sulcus in mice^26^. We therefore studied *Rnd3* gene expression in the ferret brain to ascertain if expression levels change during sulcus formation. For this, we cloned the ferret *Rnd3* cDNA and prepared an RNA probe to study its expression by in situ hybridization at early postnatal stages (P2 and P6) when brain folding starts to develop^14,15^. We noticed important differences in *Rnd3* gene expression, especially in the outer SVZ (OSVZ), at P2 and P6 (Figure 7a,b; Supplementary Figure 5). Interestingly, we observed that the ratio of *Rnd3* expression between sulcus and gyrus was significantly lower in the OSVZ compared to other layers such as the inner SVZ (ISVZ) or the CP, where expression levels were similar (Figure 7c-h), indicating that *Rnd3* expression is specifically downregulated in the sulcus OSVZ. The OSVZ has been considered an emerging new layer of progenitor cells important for brain evolution and folding^3,10,14,16,17^. Therefore, the fact that *Rnd3* is downregulated specifically in this area during sulcus formation along with our observations that *Rnd3* ablation causes spontaneous folding of the mouse brain, provides further support to the idea that *Rnd3* has an important role during cortex development and folding in vivo, in gyrecephalic species.

**Figure 7:**
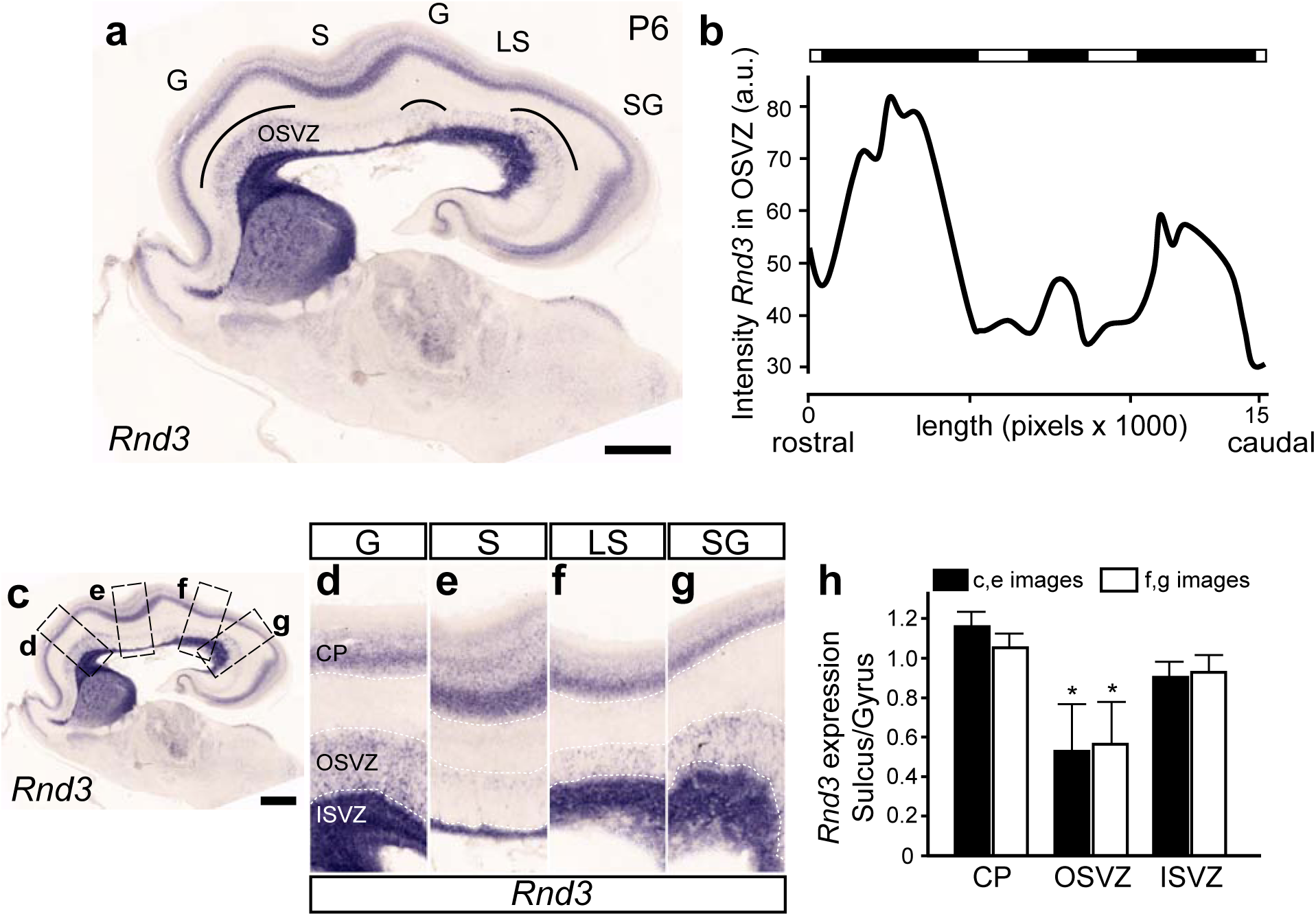
*Rnd3* is differentially expressed in the OSVZ of the developing ferret cortex. **a** In situ hybridization of *Rnd3* of sagittal sections of ferret brain at P6 (rostral left). The position of the outer subventricular zone (OSVZ), the cortical gyrus (G) and sulcus (S), including the lateral sulcus (LS) and the splenial gyrus (SG), are indicated. Curved lines in the OSVZ, indicate the areas of higher *Rnd3* expression. Scale bar, 1 mm. **b** *Rnd3* in situ hybridization signal in the OSVZ along the rostro (left)-caudal (right) length of the cortex. Horizontal bar at the top is a binary representation of *Rnd3* expression levels within the OSVZ indicating modules of overall high (black) and low (white) expression. Intensity is expressed as arbitrary units (a.u.); cortical length is expressed in pixels. **c** Overview image of *Rnd3* in situ hybridization of ferret brain at P6 (as in **a**) indicating the position of the images shown in **d**-**g** (dashed squares). Scale bar, 1 mm. **d**-**g**) High magnification of the two gyri (**d**,**g**) and the adjacent sulci (**e**,**f**), including the lateral sulcus (LS, **f**) and the splenial gyrus (SG, **g**), depicted in **c**. The cortical plate (CP), outer subventricular zone (OSVZ) and the inner subventricular zone (ISVZ) are indicated (**d**). **h** Quantification of the relative *Rnd3* in situ hybridization signal of images **d**-**g** (**d**,**e**, black bars; **f**,**g**, white bars). Values are expressed as the ratio sulcus/adjacent gyrus in the CP, OSVZ or ISVZ and as the mean±SEM; n=3 sections from 2 brains. *p<0.05, one-way ANOVA with Tukey post-hoc analysis.

## Discussion

Cortical folding is a key evolutionary trait for developing higher cognitive tasks in humans and other larger mammals. However, the molecular mechanisms that drive this crucial process are largely unknown. In this study we have revealed an important interaction between the transmembrane protein FLRT3 and the atypical Rho-GTPase RND3, during radial migration of cortical neurons and the formation of the cortical folds. Structural prediction and mutational analysis pinpoint the interaction to a β-strand-mediated binding mechanism, involving a conserved binding motif of RND. We show that both, *Rnd3* and *Flrt3*, are expressed in regions of active migration during mouse cortex development. Interestingly, reduction of RND3 leads to an increased radial migration, showing that its activity is required to regulate migration. Interestingly, genetic ablation of *Rnd3* triggers the formation of spontaneous cortical sulci in the mouse brain without affecting progenitor cell amplification, which could explain the similar cortical folding phenotype observed in *Flrt1*/*3* DKO mutants^26^. As seen for FLRTs, we found an inverse correlation of *Rnd3* expression in the OSVZ with the formation of sulci in ferret brains, favouring the idea that reduction of *Rnd3* levels is an important mechanism for sulcus formation. Our data therefore support a model in which FLRT3-RND3 negatively regulates cortical folding both in mice and in gyrencephalic species.

### Involvement of RND3 and FLRT3 in cortical folding

Most of the available models of cortical folding involve the expansion of neuron progenitor cells^63,74,75,76,77,78^. RND3 has been involved in the coordination of interkinetic nuclear migration, the regulation of the orientation of the cleavage-plane and of the adherens junctions in cortical radial glial stem cells, while in BPs, RND3 inhibits CyclinD1 and controls cell proliferation^45^. Moreover, during postnatal neurogenesis in the lateral SVZ, *Rnd3* deletion promotes neural stem cell proliferation by upregulation of Notch signaling at early postnatal stages but not at later ones^50,79^. In the present study, however, we provide evidence that progenitor expansion does not play an important role in the formation of cortical folds observed in the *Rnd3* mutant embryos: i) expression of *Rnd3* in the cortical VZ/SVZ is detected late during development, from E16.5 (Figure 1); ii) the total cell counting by DAPI staining or the number of mitotic cells (PH3+), do not show an increase in *Rnd3* KO cortices (Figure 3; Supplementary Figure 1) and iii) IUE experiments by overexpressing or deleting *Rnd3*, do not alter the total number of electroporated cells (EGFP+). These results are consistent with our previous findings where proliferation in the subpallial VZ/SVZ was not altered in the *Rnd3^KO^* embryos^49^.

Besides proliferation, neuron migration also contributes to cortex folding^14,15,18,19^. In fact, in the *Flrt1*/*3* DKO, the appearance of cortical sulci was attributed to a reduced intercellular adhesion, reflecting the role of FLRTs as homotypic or heterotypic cell adhesion molecules to Latrophilins^26,28,29^, which led to a faster and more divergent migration, without affecting proliferation^26^. In the present study we provide compelling evidence that RND3 acts as an intracellular binding partner to FLRT3 in regulating neuron migration and cortical folding, as *Rnd3* genetic ablation and the specific impairment of the RND3-FLRT3 interaction is sufficient to trigger faster migration and cortical sulci formation. Importantly, the fact that a single deletion of *Rnd3* recapitulates many of the cortical sulci features observed in the *Flrt1*/*3* DKOs^26^, strongly argues for a role of RND3 as a common signalling effector downstream of different FLRTs. In agreement, RND3 also interacts with FLRT1, although at lesser extent (data not shown), and *Flrt1* single KO also shows sulcus formation, with a lower penetrance compared to *Flrt1/3* DKO^26^. Despite we observed a faster migration of *Rnd3* deficient neurons, we were unable to detect cell clustering and tangential dispersion in the CP as in the *Flrt1/3* DKO^26^ (data not shown). This could be because *Rnd3* expression is downregulated as neurons enter the CP (Figure 1) and we cannot track the behaviour of *Rnd3* KO neurons within the CP with the LacZ reporter. Our observations in mice align well with the expression results obtained in ferret. *Rnd3* expression is specifically reduced in the OSVZ in areas of prospective sulci formation (Figure 7), indicating that *Rnd3* downregulation contributes to cortical sulci formation in gyrecencephalyc species.

### Mechanisms of regulation of cortical neuron migration by FLRT3 and RND3

The signalling mechanisms involved in the cell-adhesion properties of FLRTs are essentially unknown. In the present work we have revealed that the ICD of FLRT3 exerts an instructive signalling function through RND3 that is required for neuronal migration and the regulation of cortical folding. Based on previous findings, we propose that RND3 acts as a molecular link between FLRTs and the regulation of cytoskeleton dynamics in this context. For instance, RND3 and its relative family member RND2 were shown to regulate different steps of cortical radial migration by promoting actin cytoskeleton remodelling^36,44,80,81^. In addition, RND3 was shown to modulate migration of neuroblasts in the rostral migratory stream by disrupting cytoskeleton dynamics^49,50^. Increasing evidence suggests that RND3 regulates actin cytoskeleton dynamics by antagonizing RHOA signaling, which acts as an intracellular hub for many signaling pathways controlling cell shape and movement^82,83^. RHOA inhibition by RND3 occurs by different mechanisms during cortical neuron migration^46^ including interaction and activation of the RHOA inhibitor, p190RhoGAP^39^; blocking of RHOA effectors, such as the Rho kinase ROCKI^40^ or by inhibiting RHO guanine nucleotide exchange factors (GEFs), such as Syx^84^. While our IUE results recognize the important role of RNDs during cortical neuron migration, they differ from the proposed mechanistic model in which RNDs, by RHOA inhibition, would promote migration^36,44,81^. This might be due to important differences in the embryonic age used for IUE and the experimental approach to ablate RND expression (shRNA downregulation^44,81^ vs CRE-mediated ablation in the present study). Alternatively, these differences may be explained by the increasingly evident complexity underlying the regulation of RHOA activity by RNDs. For instance, RND overexpression, even at low levels, prevent cortical neuron migration in some circumstances^80,81^. In addition, when RHOA activity was reduced, by silencing PlexinB2, cortical neurons got stalled and overexpressing RHOA rescued their migration^52^. Accordingly, RHOA activity was reported to be necessary for the migration, proliferation, and myelin formation of Schwann cells and for the migration of cerebral granule cells or neurons in the olfactory bulb^83,85,86^. Therefore, inhibition and activation of RHOA are both necessary during neuron migration and probably reflect different needs of RHOA activity during the migratory process^44,52,83,83^. Taken together, all these studies suggest that strict regulation of RHOA activity by RNDs or other modulators is required for proper radial migration, which might be particularly important in a fast-developing structure such as the cerebral cortex. How FLRT3 regulates RND3 function? We propose the possibility that, because RND proteins are constitute active, their association with transmembrane receptors such as FLRT3 would provide a mechanism by which extracellular cues directing neuron migration can precisely control spatial regulation of RHOA activity at the plasma membrane. Consistently, we observed that the farnesylation domain of RND3 is required for cortical neuron migration, probably to facilitate the binding to membrane receptors as well as to control RHOA activity^44,87^. Alternatively, since the predicted RND3-FLRT3 binding motif is close to the amino acid responsible for p190RHOGAP binding (Thr55) ^39,44^ and overlaps with a known binding motif that engages plexin receptors (PDB code 2REX)^53,73^, which in turn compete with p190RHOGAP^52^, it is possible that FLRT3 affects RHOA activity through the modulation of p190RHOGAP or plexin binding to RND3. Future work will be necessary to ascertain this possibility. In any case, our results highlight the important role of this region as a hotspot for functional regulation of RND3 by associated partners. Finally, there is the possibility that FLRT3-RND3 interaction exerts its effects on neuron migration independently of RHOA, as RND3 was recently found to regulate lung cancer cell invasion and migration via the modulation of α5 integrin expression^88^. However, the relevance of this mechanism for cortical neuron migration needs further investigation.

### Evolutionary considerations

In contrast to the classical Rho GTPases (Rho, Rac and Cdc42), Rnd proteins emerged relatively recently during evolution and are only found in vertebrates, indicating that they might be linked to the acquisition of more specialized functions^31^. According to our results, we propose that the regulation of cortical folding could represent one such function. We show that *Rnd3* displays different expression levels in the ferret OSVZ which inversely correlates with sulci formation (Figure 7). This is consistent with the discovery of specific modules of gene expression (transcriptomic protomap) in gyrecencephalic species (but not in lisencephalic ones) which systematically map the prospective position of gyri and sulci, especially in the OSVZ^89,90^. Another important evolutionary consideration arises from the evidence that the lissencephalic mouse originates from a larger and gyrencephalic ancestor, indicating that it has undergone an evolutionary phenotypical reversal with a secondary loss of gyrencephaly^91,92,93^. In this regard, it is tempting to speculate that the FLRT/RND3 complex has had an essential role during evolution as a “break” of cortical folding and to trigger the secondary loss of gyrencephally during mammalian evolution. Therefore, it would be interesting to explore the function of these molecules in other species with secondary loss of gyrencephally such as the marmoset, which still displays a prominent OSVZ^94,95^).

## Supporting information

Cortical structure of RND3KO embryos

Expression of different RND3 constructs in HEK293T cells, co-immunoprecitation of RND3 with a FLRT3 ICD deletion mutant and effects of RND3 phosphoryl

Computational models of murine RND3 (cyan) and FLRT1-3 (green)

Sequence alignments and mutagenesis of murine RND3

Expression of Rnd3 in the P2 ferret brain

## Author contributions

B.Z., P.M-O. and J.E. conceived and designed the experiments. B.Z., P.M-O., S.A., N.N-S., M.P., Ll.V., M.B-S, V.F., D.N. and D.Ch. performed the experiments. B.Z., P.M-O., S.A., N.N-S., M.P., Ll.V., M.B-S, V.F., D.N., D.Ch. and J.E., analyzed the data. J.E. wrote the manuscript. X.D., M.E., R.K., E.S., V.B. and D.dT. improved the manuscript.

## Acknowledgements

We thank members of the Oncogenic and Developmental Signaling group for fruitful discussions. Special thanks to Sònia Rius for her outstanding technical support and to Jèssica Pairada and Leonor Gomes de Araujo for mouse colony management. We thank Dr. Daniel Sanchis (IRBLLEIDA-Universitat de Lleida, Lleida, Spain) for providing the mouse CAG-CRE line (B6.Cg-Tg(CAG-Cre)CZ-M02Osb, RBRC01828, RIKEN BRC) and Dr. Ignacio Pérez-Roger (Universidad CEU Cardenal Herrera, Valencia, Spain) for providing the *Rnd3* mouse cDNA. We thank the scientific technical facilities of the University of Lleida for their technical support.

## Supplementary Figure Legends

**Supplementary Figure 1: Cortical structure of *RND3^KO^* embryos.**

**a**-**c** DAPI staining of coronal sections of control (**a**), *Rnd3^KO^* without (w/o) sulci (**b**) or *Rnd3^KO^*with sulci (**c**) cortices at E16.5. Inner brackets indicate the cortical width measured in **d**. Scale bar, 100 μm.

**d** Quantification of the cortical width (in μm) of control (black bars), *Rnd3^KO^* without (w/o) sulci (grey bars) or *Rnd3^KO^* with sulci (white bars) brains, at E16.5, at rostral, intermediate and caudal levels. Values are expressed as the mean±SEM; n>3 sections from >2 brains/genotype. **<p=0.01, ***<p=0.001, one-way ANOVA with Tukey post-hoc analysis. No statistical differences (N.S., one-way ANOVA, p=0.058).

**e** Quantification of the total number of cells (DAPI) on a fixed area of the cortex of control (black bars), *Rnd3^KO^*without (w/o) sulci (grey bars) or *Rnd3^KO^* with sulci (white bars) brains, at E16.5, at rostral, intermediate and caudal levels. Values are expressed as the mean±SEM; n>3 sections from >2 brains/genotype. *<p0.05, one-way ANOVA with Tukey post-hoc analysis. No statistical differences (N.S., one-way ANOVA, p=0.073, intermediate; p=0.157, caudal).

**f**-**k’’’** SATB2 (turquoise), CTIP2 (red) and TBR1 (green) staining of coronal sections of control (**f**-**g’’’**), *Rnd3^KO^*without (w/o) sulci (**h**-**i’’’**) or *Rnd3^KO^* with sulci (**j**-**k’’’**) brains, at E16.5. Asterisk in **j** indicate the cortical sulci. Dashed boxes (**f**,**h**,**i**) indicate the position of panels **g**-**g’’’**, **i**-**i’’’** and **k**-**k’’’**, respectively, which show the single SATB2 (**g’’’**,**i’’’**,**k’’’**), CTIP2 (**g’’**,**i’’**,**k’’**) and TBR1 (**g’**,**i’**,**k’**) staining, including DAPI (**g**,**I**,**k**). Scale bar in **j**, 200 μm.

**Supplementary Figure 2: Expression of different RND3 constructs in HEK293T cells, co-immunoprecitation of RND3 with a FLRT3 ICD deletion mutant and effects of RND3 phosphorylation during cortical neuron migration.**

**a-e** HEK293T cells were transfected with the indicated RND3 constructs (all FLAG-tagged) including the RND3^TPM^ (phosphorylation mutant in which residues S210, S218 and S240 were substituted by alanines), RND3^ΔFS^ (deletion mutant in which the farnesylation site -FS- at the C-terminal end of the RND3 protein was deleted) and RND3^F3BM^ (RND3 mutant in which the binding motif -BM- for FLRT3 -F3- was mutated). Transfected cells were lysed for Western-blot analysis (**a**) or fixed for immunofluorescence with FLAG antibodies (**b**-**e**). In **a**, molecular weight marker is indicated on the left. In **b**-**e**, arrowheads indicate the plasma membrane localization of RND3. Scale bar, 10μm.

**f** Quantification of the percentage of transfected cells displaying RND3 localization at the plasma membrane+cytosolic clusters (grey bars), mainly at the plasma membrane (black bars) or as a diffused intracellular pattern with no plasma membrane localization (white bars). As expected, RND3^ΔFS^ displays an exclusive intracellular localization due to the lack of the CAAX motif, while RND3^TPM^ and RND3^F3BM^ are often localized at the plasma membrane. Values are expressed as the mean±SEM; n=3 experiments, >20 cells/experiment. ***p<0.001, two-way ANOVA with Tukey post-hoc analysis; N.S., no statistical difference, p=0.932.

**g**-**i** EGFP staining of coronal sections of the cortex of wild-type brains IUE with empty vector (**h**, EGFP), RND3 (**i**) or RND3^TPM^ (**j)** at E13.5 and analyzed at E19.5. The relative distribution of the electroporated cells was quantified through the cortical wall divided in ten equal bins (left). Scale bar, 100 μm.

**j** Quantification of relative distribution (%) of EGFP+ cells in lower (1-3), middle (4-7) and upper (8-10) cortical bins of wild-type embryos IUE with empty vector (EGFP, black bars), RND3 (grey bars) or RND3^TPM^ (white bars). Values are expressed as the mean±SEM; n>3 brains/condition, >5 sections/brain. *<p=0.05, **<p=0.01, one-way ANOVA with Tukey post-hoc analysis.

**k** Co-immunoprecipitation assay in transfected HEK293T cells with the indicated constructs. The lysates were immunoprecipitated with FLAG antibodies to pull down RND3 and analyzed by Western-blot with anti-FLRT3 (upper panel) or anti-FLAG antibodies (middle panel). The input levels of FLRT3 in the total cell lysates (TCL) are shown (lower panel). Molecular weight marker is indicated on the left. The bottom graph shows the quantification of the co-immunoprecipitated FLRT3. Values are expressed as fold increase respect to the non-transfected cells (1), normalized by the levels of RND3 (immunprecipitates) and FLRT3 (TCL), and as the mean±SEM; n=3 experiments).

**p<0.01, one-way ANOVA with Tukey post-hoc analysis.

**Supplementary Figure 3: Computational models of murine RND3 (cyan) and FLRT1-3 (green).**

**a** All structural models were generated using AlphaFold 3 and scores for the top solution in each run are listed.

**b** The highest ranked solutions are shown as cartoon diagrams. The sequence “fenytasf” (RND2 and 3) or “fenytacl” (RND 1) is shown in orange.

**c** Predicted FLRT interaction surfaces were calculated for the models shown in panel b, using PISA (’Protein interfaces, surfaces and assemblies’ service PISA at the European Bioinformatics Institute (http://www.ebi.ac.uk/pdbe/prot_int/pistart.html)^59^. Colours are white= not in any interface, light pink = in an interface with one of the three computed FLRT complex structures (FLRT1, 2 or 3), salmon = involved in interfaces in two of the models, red = in an interface in all three computed models. Note that the models of Rnd1 and 3 are similar to the experimentally published structure of Rnd1 (PDB 2CLS). The model of Rnd2 is disrupted in one of the beta-sheets (see arrow head) which could have resulted in the relatively fewer FLRT-interactions predicted in this area.

**d** Sequence conservation scores were calculated with Consurf^62^, using an alignment of (Mus musculus, Geotrypetes seraphini, Danio rerio, Phasianus colchicus, Pseudonaja textilis) RND 1-3. The depth of blue represents the level of sequence conservation: white= least conserved, dark blue= most conserved.

**Supplementary Figure 4: Sequence alignments and mutagenesis of murine RND3. a** Sequence alignment of murine RND protein sequences. The sequence motif highlighted in Figure 6b is highlighted.

**b** Alignment of murine FLRT protein sequences, showing the intracellular domain sequence only. The sequences interacting with RNd3 in the predicted complex models are highlighted. All alignments were done with Clustal Omega^96^.

**c** Structural models of wild type and mutant murine Rnd3 sequences were generated using AlphaFold 3. The RND3 core domain is shown. The mutated sequence (“fenytasf” in the wild type) is shown in orange and as sticks in both models. Note that the mutation causes only local structural changes according to the computational predictions. The overall protein fold appears to remain intact.

**Supplementary Figure 5: Expression of *Rnd3* in the P2 ferret brain.**

**a** In situ hybridization of *Rnd3* of sagittal sections of ferret brain at P2 (rostral left). The position of the outer subventricular zone (OSVZ) and the splenial gyrus (SG), are indicated. Curved lines in the OSVZ, indicate the areas of higher *Rnd3* expression. Scale bar, 1 mm.

**b** *Rnd3* in situ hybridization signal in the OSVZ along the rostro (left)-caudal (right) length of the cortex. Horizontal bar at the top is a binary representation of *Rnd3* expression levels within the OSVZ indicating modules of overall high (black) and low (white) expression. Intensity is expressed as arbitrary units (a.u.); cortical length is expressed in pixels.

