## Supplementary figures and images for "The atypical RHO-GTPase RND3/RHOE interacts with FLRT3 to regulate cortical migration and folding"

### Cortical structure of RND3KO embryos

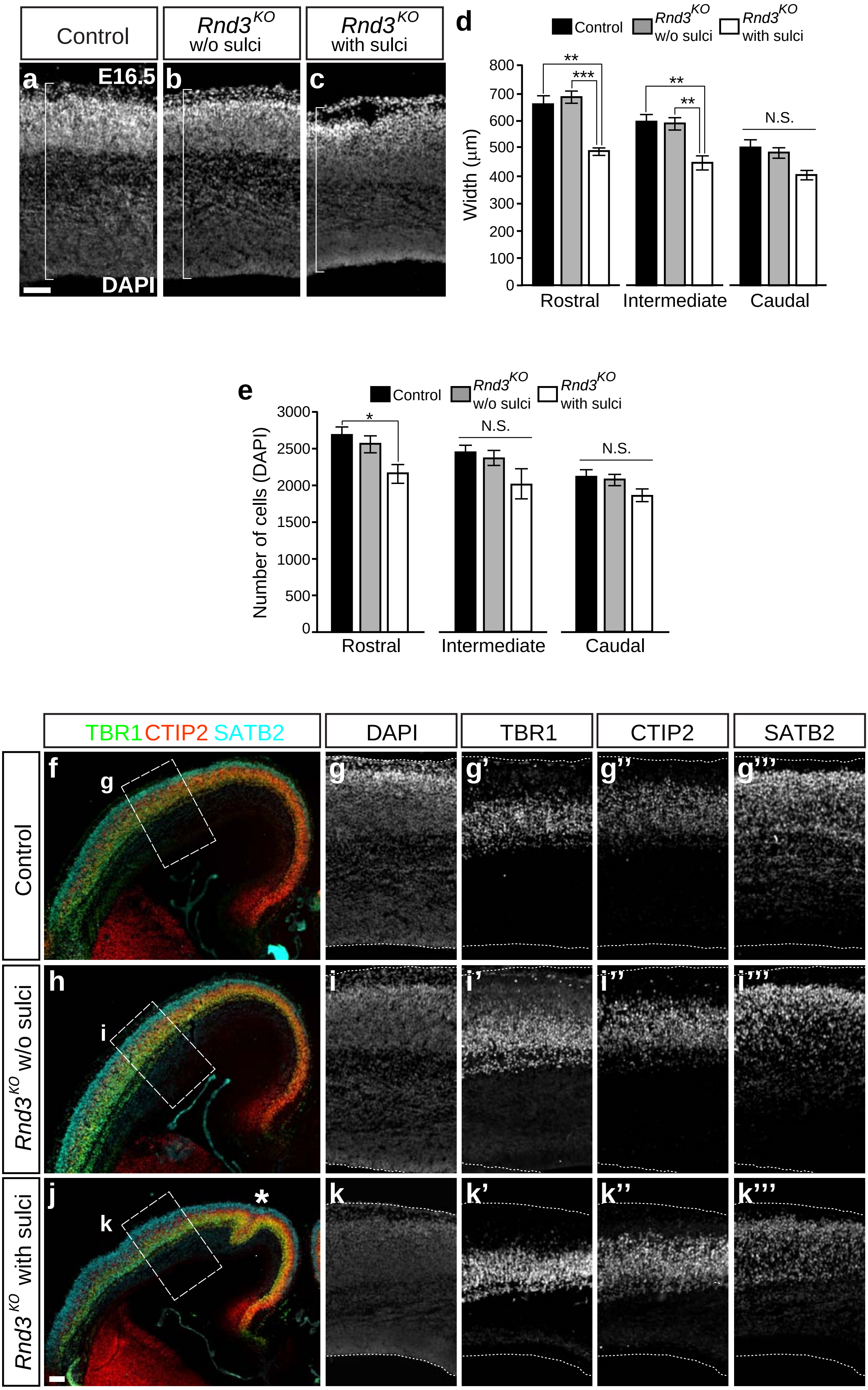


Zammou et al., Supplementary Figure 1

### Expression of different RND3 constructs in HEK293T cells, co-immunoprecitation of RND3 with a FLRT3 ICD deletion mutant and effects of RND3 phosphoryl

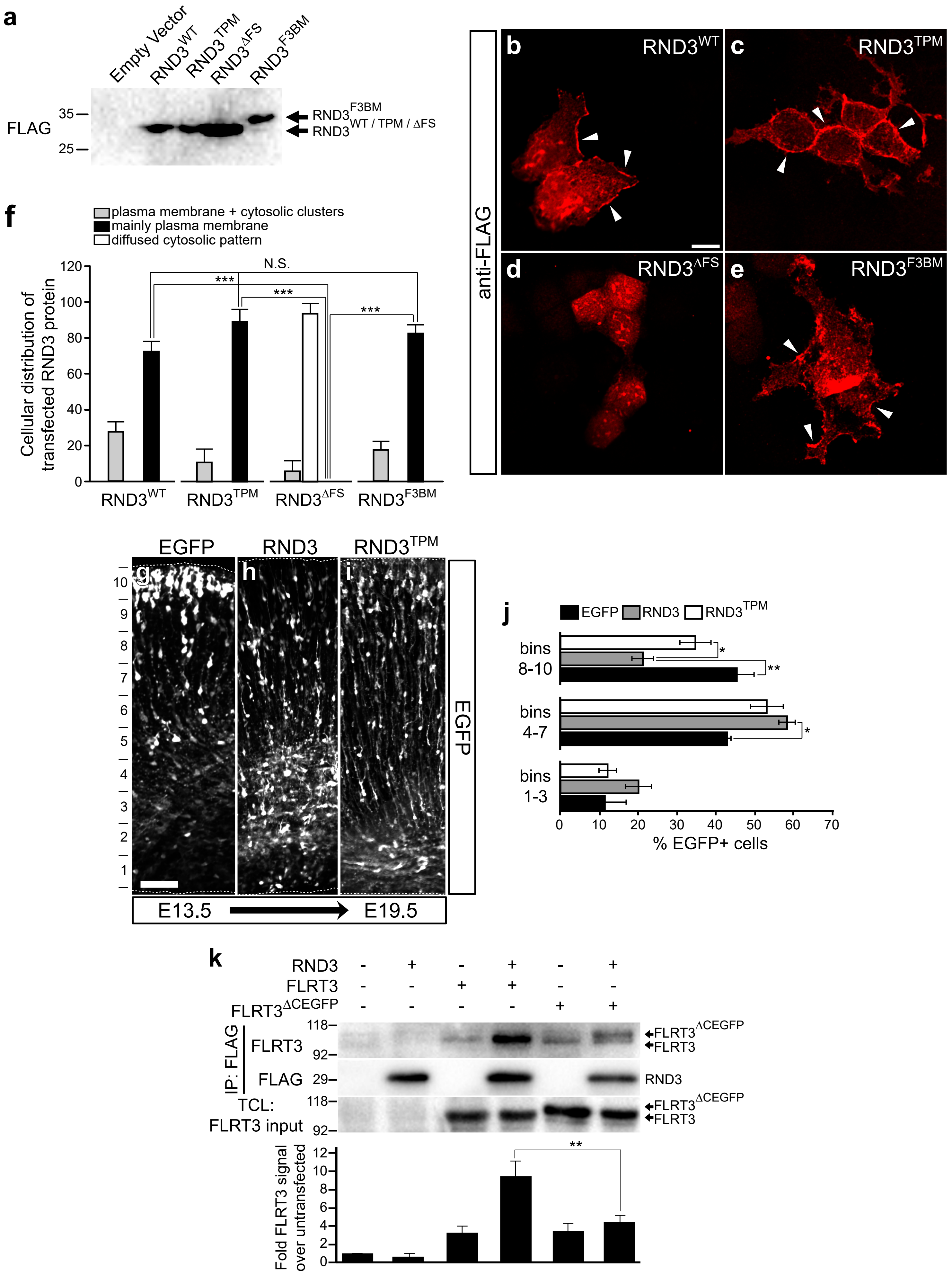


Zammou et al., Supplementary Figure 2

### Expression of Rnd3 in the P2 ferret brain

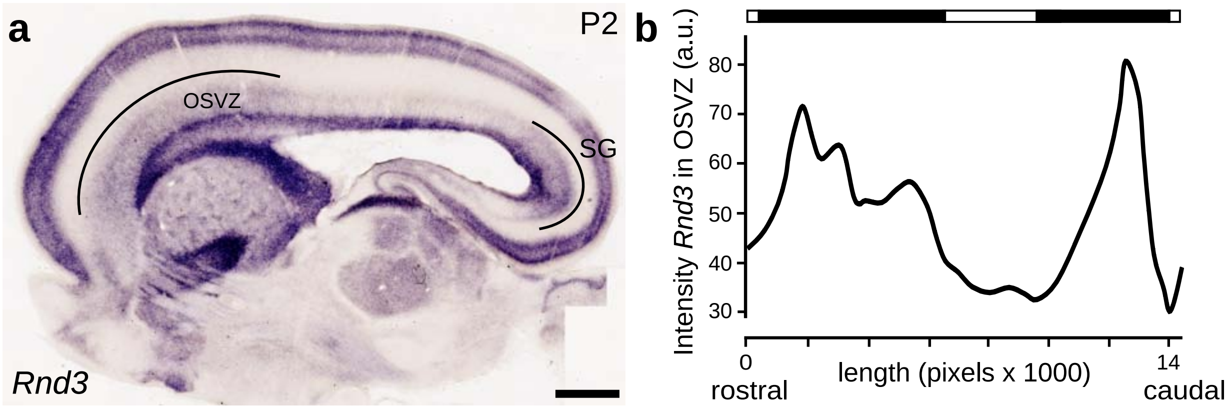


Zammou et al., Supplementary Figure 5
