## Supplementary material for "The atypical RHO-GTPase RND3/RHOE interacts with FLRT3 to regulate cortical migration and folding": Computational models of murine RND3 (cyan) and FLRT1-3 (green)

**a**

| protein complex | clash | iptm | ptm | ranking score |
|---|---|---|---|---|
| RND1-FLRT1 | no | 0.22 | 0.57 | 0.51 |
| RND2-FLRT1 | no | 0.4 | 0.58 | 0.65 |
| RND3-FLRT1 | no | 0.16 | 0.53 | 0.46 |
| RND1-FLRT2 | no | 0.26 | 0.56 | 0.54 |
| RND2-FLRT2 | no | 0.22 | 0.56 | 0.49 |
| RND3-FLRT2 | no | 0.16 | 0.53 | 0.45 |
| RND1 – FLRT3 | no | 0.23 | 0.57 | 0.52 |
| RND2 – FLRT3 | no | 0.26 | 0.57 | 0.54 |
| RND3 – FLRT3 | no | 0.16 | 0.53 | 0.46 |

**b**

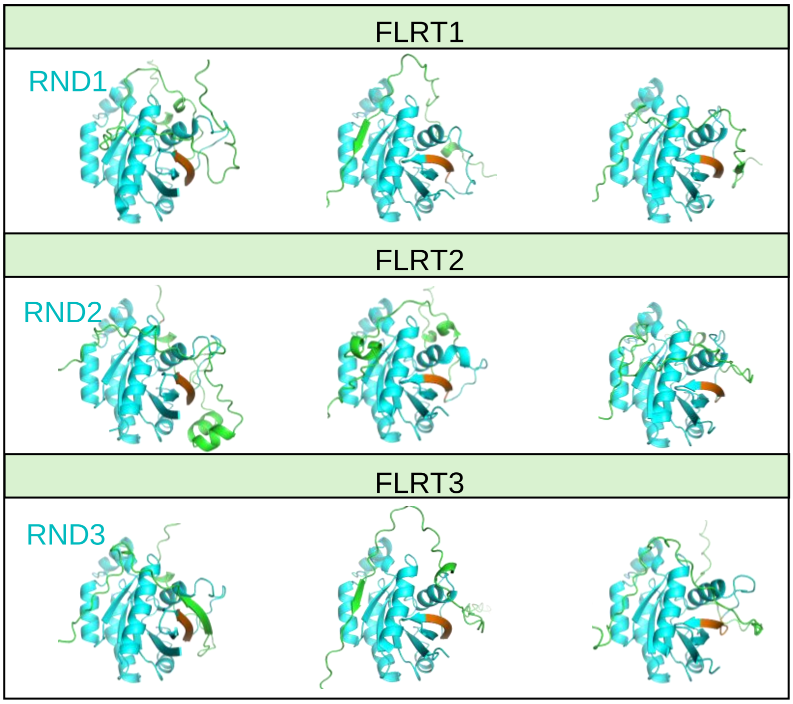


**c**

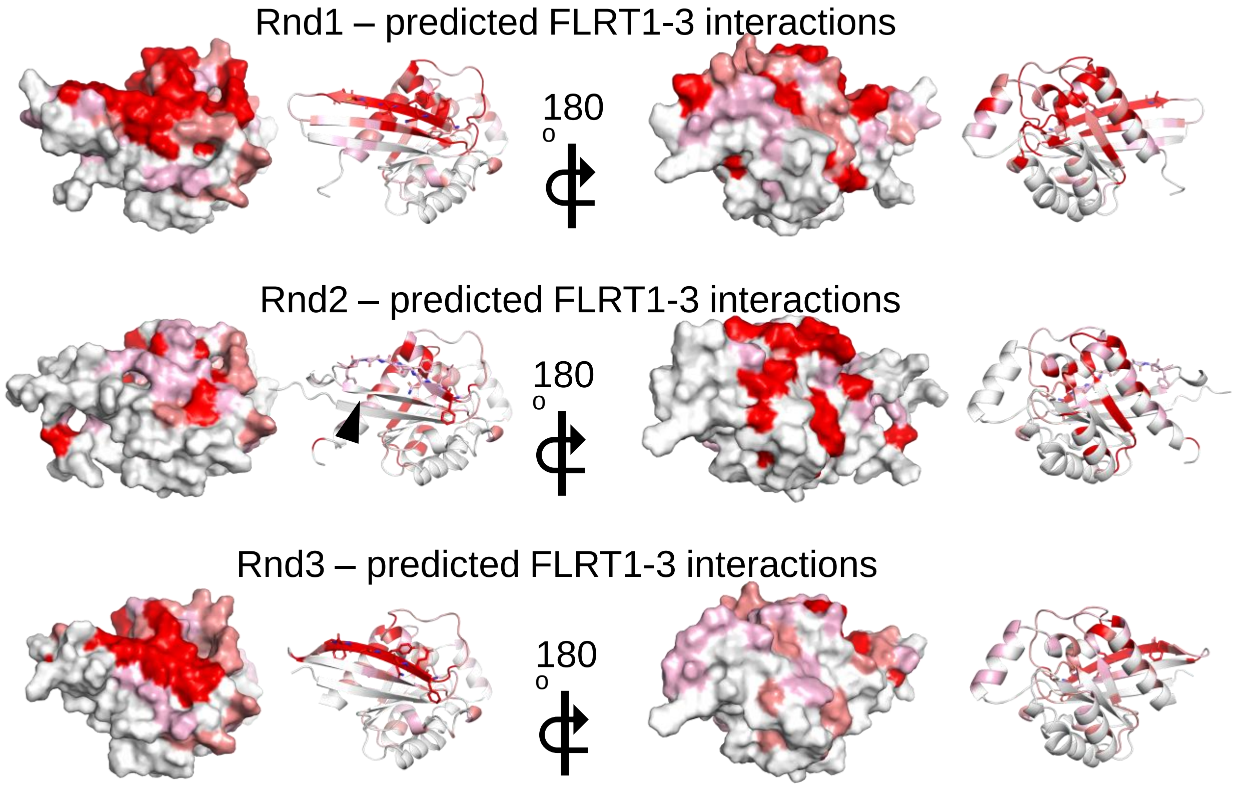


**d**

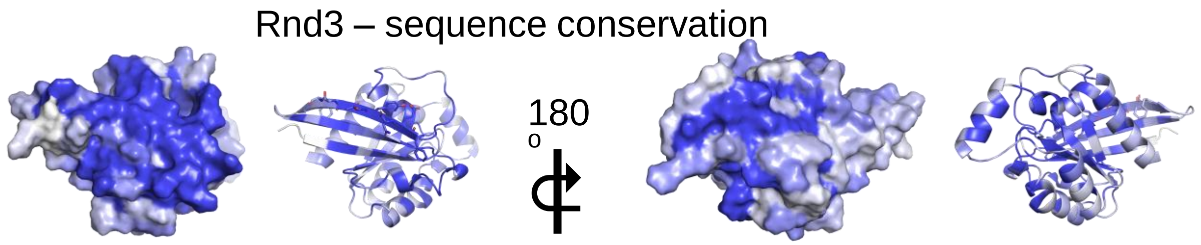


### Zammou et al., Supplementary Figure 3
