## Supplementary material for "The atypical RHO-GTPase RND3/RHOE interacts with FLRT3 to regulate cortical migration and folding": Sequence alignments and mutagenesis of murine RND3

**a**

### Murine Rnd sequences

```
Rnd1      MKERRAPQ----------PVVVRCKLVLVGDVQCGKTAMLQVLAKDCYPETYVPTVFENY  50
Rnd2      ----------------MEGQSGRCKIVVVGDAECGKTALLQVFAKDAYPGSYVPTVFENY  44
Rnd3      MKERRASQKLSSKSIMDPNQNVKCKIVVVGDSQCGKTALLHVFAKDCFPENYVPTVFENY  60
                                :**:*:*** :*****:*:*:***.:* .*********

Rnd1      TACLETEEQRVELSLWDTSGSPYYDNVRPLCYSDSDAVLLCFDISRPETMDSALKKWRTE  110
Rnd2      TASFEIDKRRIELNMWDTSGSSYYDNVRPLAYPDSDAVLICFDISRPETLDSVLKKWQGE  104
Rnd3      TASFEIDTQRIELSLWDTSGSPYYDNVRPLSYPDSDAVLICFDISRPETLDSVLKKWKGE  120
          **.:* : :*:**.:****** ********.* ******:*********:**.****: *

Rnd1      ILDYCPSTRVLLIGCKTDLRTDLSTLMELSHQKQAPISYEQGCAIAKQLGAEIYLEGSAF  170
Rnd2      TQEFCPNAKVVLVGCKLDMRTDLATLRELSKQRLIPVTHEQGTVLAKQVGAVSYVECSSR  164
Rnd3      IQEFCPNTKMLLVGCKSDLRTDVSTLVELSNHRQTPVSYDQGANMAKQIGAATYIECSAL  180
            ::**.::::*:*** *:***::** ***:::  *::::**  :***:**  *:* *: 

Rnd1      TSETSIHSIFRTASMVCLNKSSPVPPKSPVRSLSKRLLHLPSRSELI--STTFKKEKAKS  228
Rnd2      SSERSVRDVFHVATVASLGRGHRQLRRTDSRRGLQRSTQLSGRPDRGN-EGEMHKDRAKS  223
Rnd3      QSENSVRDIFHVATLACVNKTNKNVKRNKSQRATKRISHMPSRPELSAVATDLRKDKAKS  240
           ** *::.:*:.*::..:.:      :.  :   :*  :: .* :       ::*::***

Rnd1      CSIM  232
Rnd2      CNLM  227
Rnd3      CTVM  244
          *.:*
```

**b**

### Murine FLRT intracellular domain sequences

```
FLRT2     KKGRYTSQKWKYNRG-RRKDDYCEAGTKKDNSILEMTETSFQIVSLNNDQLLKGDFRLQP
FLRT1     RAGELLTRERVYNRGSRRKDDYMESGTKKDNSILEIRGPGLQMLPINP-YRSKEEYVVHT
FLRT3     RNGSLFSRNCAYSKGRRRKDDYAEAGTKKDNSILEIRETSFQMLPISNEPISKEEFVIHT
          : *   :::  *.:* ****** *:**********:   .:*:: :.     * :: ::

FLRT2     IYTPNGGINYTDCHI---PNNMRYCNSSVPDLEHCHT
FLRT1     IFPSNGSSLCKGAHTIGYGTTRGYREAGIPDVDYSYT
FLRT3     IFPPNGMNLYKNNLS-ESSSNRSYRDSGIPDSDHSHS
          *:  **    ..        ..  * ::.:** ::.::
```

**c**

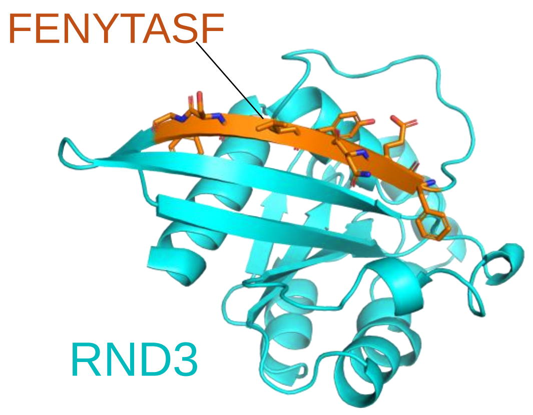


**d**

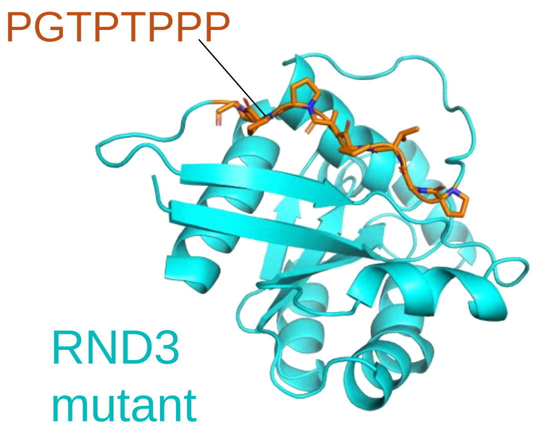


Zammou et al., Supplementary Figure 4
